# DNA Twist at High Alkaline Ion Concentrations: Evidence against C-Form DNA in Solution

**DOI:** 10.1101/2025.09.15.676196

**Authors:** Koen R. Storm, Christian Wiebeler, Sergio Cruz-León, Nadine Schwierz, Jan Lipfert

## Abstract

DNA is highly negatively charged, making its structure strongly dependent on the ionic environment. DNA twist –a central DNA property– varies with ion concentration and identity. Prior studies have focused on salt concentrations below 1 M and it is unclear whether twist trends persist at higher concentrations. It has been proposed that at high salt, DNA transitions from its canonical B-form to C-form, originally observed by fiber diffraction.

Here, we use single-molecule magnetic tweezers to measure DNA twist in high concentrations of LiCl, NaCl, KCl, and CsCl. For all salts, twist initially increases approximately as ∼[salt]^1/2^, but plateaus above 3 M. LiCl causes the largest twist increase, by ≤ 0.8°/bp compared to physiological salt, still far below the suggested C-form values of 2-3°/bp. We perform all-atom molecular dynamics simulations for DNA in LiCl solutions with different force fields. For parmbsc1, we observe good agreement with experiments when ion activities are taken into account. We find that simulations initiated in the C-form rapidly convert to B-form, while the B-form remains stable. Our results demonstrate ion-specific, systematic changes in DNA twist beyond 1 M salt, but do not support a transition to the C-form for DNA, even at very high salt concentrations.

## INTRODUCTION

DNA is the storage medium of all cellular life and increasingly used as a construction material in bionanotechnology. Since DNA is highly negatively charged, many of its properties sensitively depend on the ionic environment (1). In particular the twist of the DNA helix, famously discovered by Watson and Crick (2), depends on the concentrations and type of ions in solution (3–6). Initial measurements using a bulk assay with plasmid DNA found that DNA twist increases with increasing ion concentration and observed an approximately logarithmic dependence on ion concentration for monovalent ions in the range 50-300 mM (3,4). More recent measurements using single-molecule magnetic tweezers probed the changes in DNA twist up to a concentration of 1 M for all alkaline ions (5–7). The data reveal increasing DNA twist with increasing concentration following a power law dependence. While simple electrostatic theories predict the saturation of ionic effects at concentrations around 1 M monovalent salt (1), it is currently unclear whether and how the changes in twist continue above 1 M monovalent salt. Understanding DNA under conditions of high salt and low water activity (8) critically test models of nucleic acid structure and of DNA-ion interactions, which have typically been developed and tested only for lower concentrations. In addition, such conditions occur in halophiles, i.e. organisms that are exposed to extremely high salt conditions (9–12), such as *Halobacterium salinarum*, which grows best in 3-5 M monovalent salt (9). In addition, high salt concentrations and ion specific effects have been suggested to play critical roles in early life on earth (13–16).

In addition to ion concentration, ion identity also plays an important role. For Na^+^, K^+^, Rb^+^, and Cs^+^ the concentration dependencies of DNA twist follow very similar trends and the twist at a given concentration increases with ion size. In contrast, Li^+^ exhibits a stronger dependence on ion concentration than the other alkaline ions and induces the largest absolute twist of all alkaline ions. Molecular dynamics (MD) simulations complement these observations by providing atomic-level insight into DNA structure and dynamics. For example, they elucidate specific ion–DNA binding patterns, including the strong affinity of Li□ to the phosphate groups of the DNA backbone, which underlies its pronounced effect on DNA twist and stiffness (5,6,17,18). However, the extent to which ion–DNA interactions continue beyond 1 M and affect DNA conformations is still not well understood.

Early work based on X-ray fiber diffraction suggested the existence of different conformational forms of double-stranded DNA, termed A-, B-, and C-form (19–21). While the B-form is the accepted canonical form of double-stranded DNA under typical solution conditions, the C-form was first observed in Li-salts of DNA fibers (20,22) and later reported more broadly in fibers under conditions of high salt and low humidity (23,24). The C-form of DNA has been considered a distorted version of the B-form with a twist per base pair about 2-3° larger than the B-form (25,26). The C-form of DNA has recently been suggested to play a role in homologous recombination (27) and for certain sequences involved in strong bending (28).

In MD simulations, B- and C-form DNA can, in addition to the characterization by twist, be distinguished by the relative populations of two backbone conformations known as BI and BII, which reflect different torsional states in the DNA backbone. B-DNA is characterized by a predominance of the BI state, whereas an increased population of the BII conformation is associated with structural features observed in C-form DNA, including the higher helical twist (28,29).

Work on DNA using circular dichroism (CD) spectroscopy has suggested changes of the DNA helix at very high salt concentrations, up to ≈10 M monovalent concentration (30,31). The observed changes in the CD spectra were attributed to the canonical B-from helix transitioning to C-form in solution (30–33). Similar to the magnetic tweezers-based twist measurements at lower salt, Li^+^ was found to have the largest effect on the observed spectra of all monovalent ions investigated. However, the interpretation of the CD spectroscopy results has subsequently been challenged (34,35) and has recently been revisited with additional measurements on circular DNA (36). Taken together, the evidence for C-DNA in solution remains controversial and it is an open question to what extent the high twist C-form conformations observed under low humidity in fiber diffraction data are indeed also populated in solution.

Simulations have also begun to address conformational shifts and B- to C-DNA transitions (28). Several force fields including CHARMM36 (37), the AMBER-based force fields OL21 (38), parmbsc1 (39), and TUMUC1 (29) improve DNA backbone accuracy by refining torsional parameters and enhancing the BI/BII equilibrium. However, the question of C-DNA formation under high-salt conditions remains largely unexplored in MD simulations.

Here, we use single-molecule magnetic tweezers measurements (**Figure 1a**) to systematically monitor changes in DNA for high alkaline ion concentrations, Li^+^ in particular, up to 8 M. The results suggest that DNA twist increases with increasing Na^+^, K^+^, Cs^+^ and Li^+^ concentrations up to 4-5 M, i.e., well beyond 1 M that was previously investigated. At even higher Li^+^ concentration the DNA twist saturates, which challenges the previous claims of C-DNA formation at high LiCl concentrations. Complementary measurements using CD spectroscopy reveal change in the spectra fully consistent with previous experiments and indicative of structural changes of the DNA helix even beyond 5 M LiCl. Our all-atom MD simulations in LiCl support the experimental findings and reproduce the observed twist trends, in particular when ion activity is considered. The simulations further indicate that B-form DNA remains stable under high salt, while C-form conformations are not maintained. Together, these approaches provide a comprehensive view of DNA behavior in high salt conditions and challenge longstanding assumptions about high-salt-induced structural transitions.

**Figure 1.**
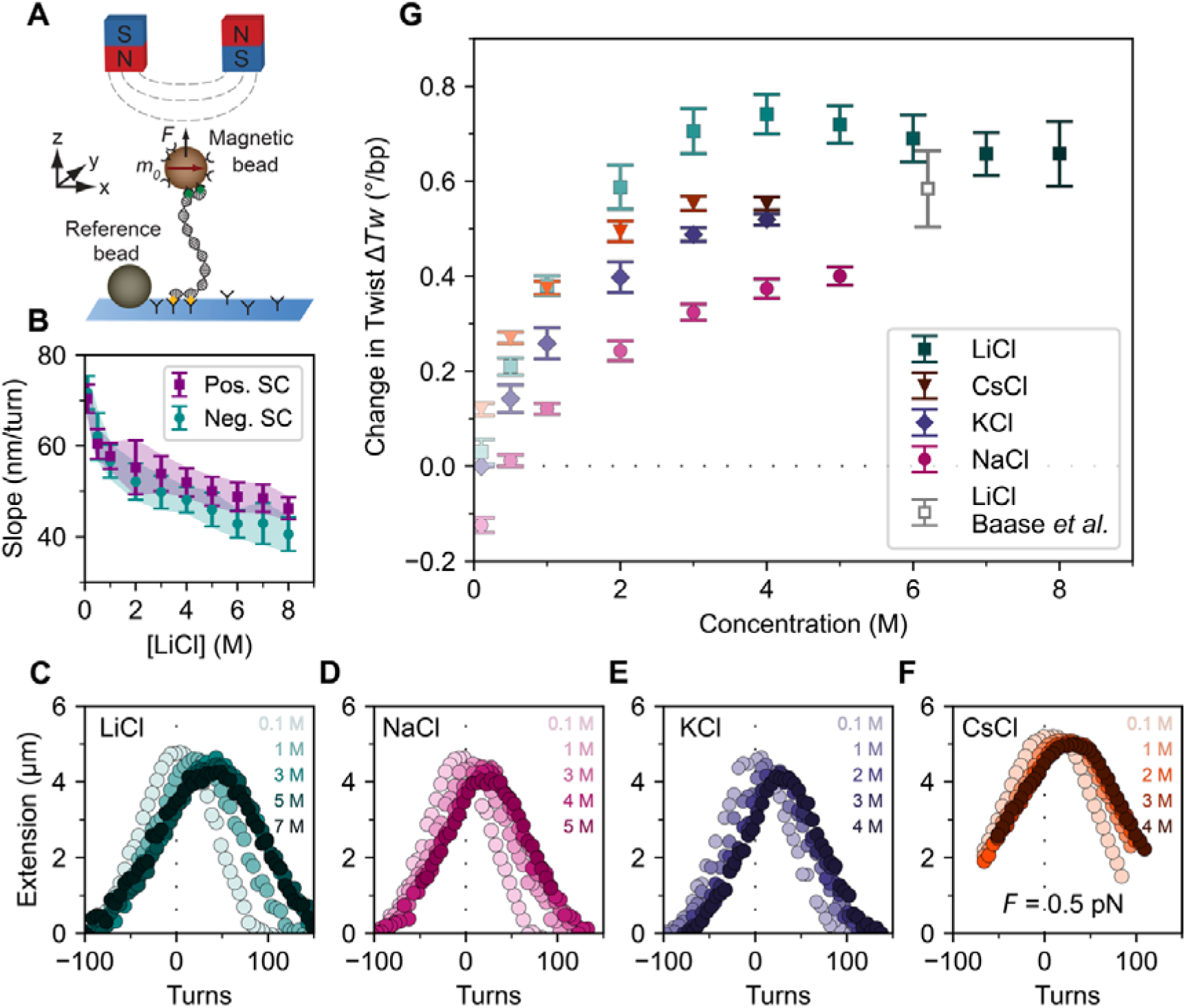
Magnetic tweezers probe changes in DNA twist with increasing LiCl concentration. A) Schematic of the magnetic tweezers setup. A DNA molecule is tethered between the flow cell surface and a magnetic bead via multiple attachment point at both ends. External magnets enable applications of forces and torques to the DNA tether. Reference beads are tracked for drift correction. B) Extension vs. turns slopes in the plectonemic regime from fits to the rotation curve data (including the data in panel C) at *F* = 0.25 pN as a function of LiCl concentration for positive (purple squares) and negative (cyan circles) plectonemic supercoils. Data points and error bars are the mean ± SD from at least 11 molecules. Similar data for other ions are shown in **Supplementary Figure S9**) DNA extension vs. applied rotation measurement in the presence of increasing concentrations of C) LiCl, D) NaCl, E) KCl and F) CsCl. Zero turns is defined as the center of the extension vs. rotation curves in 100 mM KCl, i.e. the linking number at which the DNA torsionally relaxed in the reference conditions of 100 mM KCl. The curves shift systematically to larger turns with increasing salt concentration, indicative of increasing helical twist with increasing concentration. Data are at *F =* 0.25 pN, except for CsCl, which was recorded at 0.5 pN since CsCl has a tendency to collapse DNA (**Supplementary Figure S7**). G) Change in DNA twist determined from the centers of the rotation curves (including the data in panels C-F, **Supplementary Figure S3**) as a function of monovalent salt concentration. Data are relative to 100 mM KCl and represent the mean ± SD from at least 3 and typically 10-20 molecules. The grey data point was obtained by analytical ultracentrifugation and taken from Ref. (34).

## MATERIALS AND METHODS

### Chemicals

1× phosphate buffered saline (PBS, 10 mM phosphate buffer, pH 7.4, 138 mM NaCl, 2.7 mM KCl; Sigma-Aldrich), prepared with ultrapure water (Milli-Q; Millipore), was used for DNA incubation. 10 mM Tris (Pre-set, pH 7.0, pHast Pack; Sigma-Aldrich), prepared with ultrapure water (Milli-Q; Millipore), was used for salt solution preparation. 0.1-5 M NaCl (99.5 %, p.a., ACS, ISO; Carl Roth), 0.1-4 M KCl (99.5 %, p.a., ACS, ISO; Carl Roth), 0.1-12 M LiCl (99 %, p.a., ACS; Carl Roth) and 0.1-7 M CsCl (99.999 %, p.a., Ultra Quality; Carl Roth) were prepared using 10 mM Tris (pH 7.0) at room temperature (≈22 °C) in a volumetric flask (25 ± 0.04 mL; DURAN) and stored in glass vials (Sigma-Aldrich). Salts were protected against humidity. 100 mM KCl was used as reference condition for MT measurements, matching our previous choice of reference conditition (5). Refractive indices were determined (λ=589.3 nm, 21 °C; Abbe Refractometer 3T, Atago) to confirm the salt concentration and for MT analysis (**Supplementary Figures 1 and 2 and Supplementary Table 1**). Our results for the refractive indices are in excellent agreement with literature values (40) and expand the range previously covered for LiCl.

### DNA construct and flow cell preparation

Our magnetic tweezers measurements used a 20.6 kbp DNA construct, based on the λ-phage DNA sequence and prepared as described previously (41,42). The DNA construct is designed to allow torsionally constrained attachment to the bottom glass surface of a flow cell and paramagnetic beads, respectively. The sequence of our DNA construct is representative of “random”, genomic DNA and provides effectively a sequence averaged view of DNA’s response, as is typical for MT measurements. In brief, PCR-generated DNA fragments (≈ 600 bp) labeled with multiple biotin or digoxigenin groups were ligated to the DNA (20.6 kbp), to bind streptavidin-coated magnetic beads (1.0 μm diameter MyOne beads, Thermo Fisher Scientific Baltics UAB, LT) and the flow cell surface, respectively. To attach the DNA to the flow cell, the bottom coverslip was first modified with (3-Glycidoxypropyl)trimethoxysilane. Next, 100 μL of a 5000x diluted stock solution of polystyrene beads (Polysciences, USA) in ethanol (Carl Roth, Germany) was dropcasted on the silanized slides, left to dry while covered. The polystyrene beads serve as fiducial markers for drift correction. A laser cutter was used to cut openings with a radius of 1 mm in the top coverslip for buffer exchange. The two coverslips were glued together by a single layer of melted Parafilm (Carl Roth, Germany), forming a ≈50 μL channel that connects the inlet and outlet opening of the flow cell. After flow cell assembly, 100 μg/mL anti-digoxigenin (Abcam plc., UK) in 1× PBS was introduced and incubated for at least 1 h. To reduce non-specific interactions of the DNA and beads with the flow cell surface, the flow cell was flushed with 100 μL of 250 mg/mL bovine serum albumin (BSA; Carl Roth, Germany), incubated for 1 h and rinsed with 600 μL of 1× PBS. Next, the DNA construct was attached to streptavidin coated beads by incubating 1 μL of picomolar DNA stock solution and 2 μL MyOne beads in 200 μL 1× PBS (Sigma-Aldrich, USA) for 5 min and subsequently introduced in the flow cell for 20 min to allow formation of digoxigenin-anti-digoxigenin bonds. Subsequently, the flow cell was rinsed with 2 mL of 1× PBS to flush out unbound beads.

### Magnetic tweezers instrument

Multiplexed single molecule magnetic tweezers experiments were performed as previously described (5,43,44). For twist control and the application of precisely calibrated forces we employ a pair of permanent magnets (5 x 5 x 5 mm^3^, Supermagnete, Switzerland) arranged in vertical geometry (45) with a gap of 1 mm and mounted in a holder that is controlled by a translation motor (M-126.PD2, PI) and a rotation motor (C-150.PD, PI). The motors are controlled, and magnetic beads are tracked in real-time by processing the camera images using a custom-written LabView program (46,47). An LED (69647; Lumitronix LED Technik GmbH) provides illumination. The field of view is imaged using a 40× oil-immersion objective (UPLFLN 40×; Olympus), together with a CMOS sensor camera (12M Falcon2; Tele-dyne DALSA). A frame grabber (PCIe 1433; National Instruments) collects images recorded at 58 Hz. A look-up table is created by moving the objective with on a piezo stage (PifocP726. 1CD, PI Physikinstrumente). Position measurements were corrected for the change in refractive index with salt concentration (**Supplementary Figure S2**).

### Magnetic tweezers twist measurements

Suitable double-stranded DNA tethers were identified by rotating the external magnets to introduce negative supercoiling under high tension (*F* ≥ 5 pN), where for single DNA tethers the formation of plectonemes at negative linking differences is suppressed due to melting and the extension remains unchanged (48). In contrast, if multiple tethers are attached to the same bead, negative supercoiling results in braiding, which decreases tether extension. To assess whether DNA tethers were torsionally constrained, positive linking differences are introduced at low force (*F* = 0.5 pN), which results in plectoneme formation and a corresponding decreasing DNA extension for torsionally constrained tethers. In nicked DNA tethers, no linking difference can be introduced, and the extension remains constant on magnet rotation. Beads bound by multiple tethers or nicked tethers are discarded from further analysis. Following bead selection and testing, the buffer in the flow cell was exchanged for a buffer comprising 100 mM KCl and 10 mM Tris (pH = 7.0), since this is the reference condition employed previously (5). We recorded extension-rotation curves by changing the magnet rotation, and thus the linking number ΔLk, in steps of five turns at low force (*F* = 0.25 pN unless stated otherwise). At each step of five applied turns, the extension of the tether is recorded for 30 s (at a camera rate of 58 Hz), and the mean extension is calculated. Subsequently, the buffer was exchanged to vary salt conditions. All measurements used 10 mM Tris, pH = 7.0, as buffer and were performed at room temperature, 22 °C. To ensure equilibration, we flushed a large volume of buffer at each new concentration through the flow cell (>300 μL or ≈6 cell volumes). During buffer exchange, we apply high force *F* = 2 pN, to ensure that the tethered beads do not rotate under the liquid flow and maintain constant linking number.

### Circular dichroism measurements

CD spectra were collected using a spectropolarimeter (J-810-150S, JASCO, Japan) equipped with a temperature controller (20 °C; CDF-426S, JASCO, Japan) in 10 × 10 mm sealable quartz cells (Hellma Analytics). Samples were prepared using the same stock for buffer and salt solution used for MT measurements (see above), i.e., 12 M LiCl (LiCl; >= 99 %, ACS; Carl Roth), but employed 40 μg/mL λ-DNA (Lambda DNA, 0.3 μg/μL, 48.5 kbp, 10 mM Tris-HCl (pH 7.6) and 1 mM EDTA; catalogue number SD0011 from Thermo Scientific) with a final volume of 2500 μL. We employed λ-DNA, instead of the functionalized DNA construct used in MT measurements, since the functionalized DNA is difficult and wasteful to produce in sufficient quantities for CD spectroscopy and since the labels used for attachment in the MT might interfere in the spectroscopic measurements. Measurements were performed with a scan rate of 20 nm/min and spectra were obtained as the accumulation of three scans. The individual DNA concentrations were determined from the absorbance using a molar absorptivity at 260 nm of 6600 M^-1^ cm^-1^. CD was expressed as Δε = ε_L_ *–* ε_R_ in units of M^-1^ cm^-1^. The molarity (M) was related to mean residual weight in the DNA samples, i.e. to the nucleotide concentration, which is independent of DNA length.

### Molecular dynamics simulations

We performed all-atom molecular dynamics simulations of a 33 bp double-stranded DNA, in each case comparing simulations using the B- and the C-form as starting structures. If not stated otherwise, the Gromacs software package in version 2020.7 (49) was employed for system preparation and simulation. For electrostatics, the particle-mesh Ewald (PME) summation was used with an order of 4 and a Fourier space grid of 0.12 nm. Furthermore, a cut-off of 1.2 nm was employed for Lennard-Jones and close Coulomb real space interactions. A time step of 2 fs was used and the LINCS algorithm was employed to constrain hydrogen bonds.

To generate structural models, we used the same DNA sequence as used previously (5) and employed Web 3DNA 2.0 (50) to generate starting structures based on fiber models in B- and C-form DNA geometry, respectively (ID 4 and 7 in the nomenclature of Ref. (25)). The resulting B-form structure was in close agreement with our previous model with a root mean square displacement (RMSD) of 1.0 Å for the backbone atoms, whereas the RMSD between the models for B- and C-Form DNA was 4.1 Å. One prominent difference between the two generated structures is the helical twist, which is 38.7°/bp for the C-form, compared to 36.1°/bp for the B-form. In our simulations, we tested several DNA force fields in combination with specific ion and water models. As one combination, which was also employed in previous work (5), we used the parmbsc1 force field (39) for DNA, combined with Mamatkulov–Schwierz ion parameters (51), and TIP3P water (52), across a broad range of ion concentrations. The choice of the ion force fields was motivated by the fact that the parameters were optimized to yield accurate ion–water and ion–ion interactions as judged by the comparison to experimental solvation free energies and activity coefficients up to 3 M salt concentration (51). For the larger salt concentrations, we computed the radial distribution functions, determined the activity derivatives from Kirkwood-Buff theory (53,54), and compared to experimental data (**Supplementary Methods**). Even at the largest salt concentration there was no indication of clustering or crystal formation.

To set the ion concentration in the simulations, we initialized simulations at defined molalities (i.e. moles of ions per kg water). Comparison to the experimental data however required concentration in molarity (i.e. moles of ions per liter of water). Therefore, we calculated the box volumes, by averaging the volume over the time of the production simulations in the NPT ensemble. In **Supplementary Tables S3 and S4**, we report the concentrations in molality and in molarity and the volumes for the parmbsc1 simulations used in this work. Throughout the work, we report the concentrations in molarity, since this is what is controlled experimentally, unless otherwise noted.

We also carried out simulations using the CHARMM36 force field (37) for DNA, while maintaining the same ion and water parameters, for eight LiCl concentrations from 1 to 6.5 M (1 m to 8 m on the molality scale). In addition, we tested the OL21 force field (38) for DNA, again with the same ion and water models, at two LiCl concentrations: 1 M and 4.4 M (1 m and 5 m on the molality scale). Similarly, simulations using the TUMUC1 force field (29) were performed at 1 M and 4.4 M LiCl with identical ion and water settings. Finally, to reproduce the conditions used by Strelnikov et al. (28), we conducted simulations using parmbsc1 for DNA, Joung–Cheatham ion parameters (55), and TIP4P-Ew water (56), at 1 M and 4.5 M LiCl. The double-stranded DNA structures in B- or C-form were put in a rhombic dodecahedron box with a minimal distance of 1.5 nm to the edges and filled with water molecules.

For all systems, the initial simulation protocol consisted of energy minimization followed by NVT and NPT simulations. During this protocol, the heavy atoms of the nucleic acids were restrained (1000 kJ/(mol nm^2^)) to allow solvent equilibration. Energy minimization with the steepest descent algorithm used a maximum of 50,000 steps. Subsequently, we used NVT and NPT simulations of 1 ns each for obtaining systems at ambient conditions. In these simulations, the temperature of 300 K was maintained using the velocity rescaling thermostat (57) with a coupling constant of 0.1 ps. In the NPT simulations, the isotropic Parrinello-Rahman barostat (58) with a coupling constant of 5 ps was employed in addition.

After this initial protocol, we first performed 300 ns simulations in the NPT ensemble without restraints. Starting from the final equilibrated structures, three 300 ns production runs were carried out, and the first 100 ns of each trajectory were discarded from the analyses. The simulation volumes determined from NPT ensemble simulations (**Supplementary Tables S3 and S4**) fluctuate by less than 0.2% (or ≤ 3 nm^3^) in the course of the trajectories. The reported volumes include the volume of the DNA, which is 2% (≈ 40 nm^3^) of the total simulation volume.

The helical properties were determined with the 3DNA software (25,59) by using the do_x3dna software package together with its dnaMD python module (60) and in-house scripts. To determine the number of adsorbed ions, we counted for each frame all Li^+^ that were within a cutoff distance of 0.3 nm to any non-bridging backbone oxygen atom of the double-stranded DNA and averaged this over each trajectory. For each concentration and DNA form, averages and standard errors of the mean were estimated from the three independent production runs. Values reported in our previous study (5) were employed for determining changes of helical properties relative to 0.10 M KCl and for the results up to a maximum concentration of 1 M.

For obtaining representative structures, we employed clustering with the algorithm of Daura et al. (61). Structures were taken from all three production runs per concentration and initial B- or C-form. A cutoff of 0.5 nm was employed, to group similar conformations based on RMSD. In each case, the largest cluster encompassed at least 90 % of all structures and the central structure of this cluster is therefore considered to be representative.

Gromaps (62) was used to analyze the three-dimensional cation distribution, and the results were visualized with pymol (The PyMOL Molecular Graphics System, Version 3.1 Schrödinger, LLC, New York, NY, USA).

## RESULTS AND DISCUSSION

### Single-molecule magnetic tweezers measurements determine DNA twist for high salt concentrations

We used single-molecule magnetic tweezers (MT) to manipulate 20.6 kbp DNA molecules tethered between a functionalized flow cell surface and small, 1.0 µm diameter, superparamagnetic beads (**Figure 1A** and Materials and Methods). The MT allow to apply precisely calibrated stretching forces and to control the rotation of the magnetic bead and thus the linking number of the tethered molecules (45,63–66). We systematically measured the DNA tether extension as a function of applied turns under a low stretching force (**Figure 1C-F**). In this low force regime, the force response of DNA is dominated by entropic stretching and well-described by the inextensible worm-like chain model (67,68). The response of the tether extension to applied turns is approximately symmetric for over- and underwinding of the helix and the DNA undergoes a buckling transition for both positive and negative applied turns (63,69). Past the buckling transition, the DNA forms positive or negative plectonemic supercoils, respectively, marked by a linear decrease of the tether extension with the number of applied turns (**Figure 1B-F and Supplementary Figure S3**). The rotation-extension curves reveal systematic shifts to positive turns with increasing ion concentration (**Figure 1**). We quantify the shifts by fitting the linear slopes in the plectonemic regime and determining the intersection points of the slope at positive and negative turns (5,70) (**Supplementary Figure S3**). Importantly, the changes are fully reversible: if the salt concentration is lowered again, the curves return to their previous position (**Supplementary Figure S4**).

To facilitate direct comparison to previous measurements (5), we used 100 mM KCl as our reference conditions. This choice of reference condition was motivated by the considerations that i) 100 mM monovalent ions correspond, approximately, to physiological ionic strength and intracellularly K^+^ is the most abundant cation, ii) 100 mM KCl is a condition that was included in previous studies of the salt dependence of DNA twist (3,4) and iii) 100 mM KCl is readily and robustly accessible both in MT experiments and MD simulations.

### DNA twist increases up to 4-5 M monovalent ion concentration

We find that DNA twist increases with increasing LiCl, NaCl, KCl, and CsCl concentration, consistent with previous measurements (**Figure 1G**). Our data up to 1 M are in good agreement with previous measurements that used a very similar methodology, but a different DNA construct, of a different length and DNA sequence (5) (**Supplementary Figures S5**). Interestingly, we observe a significant increase of DNA twist when increasing ion concentration beyond 1 M. For all tested ions, LiCl, NaCl, KCl, and CsCl, the twist increases steadily up to 3-4 M. For NaCl and KCl, we continued measurements until the maximum aqueous solubility (4 M for KCl and 5 M for NaCl), reaching a change in twist of 0.4 and 0.5°/bp, respectively (**Figure 1G**), relative to the reference conditions of 100 mM KCl.

LiCl induces the largest DNA twist of all the ions tested. Similar to the other ions, the twist increases for Li^+^ concentrations beyond 1 M: from ≈0.4 °/bp at 1 M to ≈0.74 °/bp at 4 M, all again with respect to the 100 mM KCl condition (**Figure 1G**, **Table 1 and Supplementary Figure S6**). Beyond 4 M Li^+^, the rotation-extension curves do not shift further along the turns-axis, suggesting that the average twist of DNA helix remains approximately constant and even slightly decreases (though values remain within experimental error) upon further increasing the concentration (**Figure 1G**). For LiCl, concentrations above 8 M resulted in detachment of the DNA constructs in the flow cell, preventing measurements up to the solubility limit of LiCl (≈ 14 M).

**Table 1.** Change in DNA twist (relative to 100 mM KCl) induced by alkaline ions. Data are the mean ± standard deviation from at least 3 and typically 10-20 measurements.

| Concentration<br>(M) | LiCl<br>$\Delta T_w$ (°/bp) | NaCl<br>$\Delta T_w$ (°/bp) | KCl<br>$\Delta T_w$ (°/bp) | CsCl<br>$\Delta T_w$ (°/bp) |
| --- | --- | --- | --- | --- |
| 0.1 | $0.03 \pm 0.03$ | $-0.12 \pm 0.02$ | 0 | $0.12 \pm 0.01$ |
| 0.5 | $0.21 \pm 0.02$ | $0.01 \pm 0.01$ | $0.14 \pm 0.03$ | $0.27 \pm 0.01$ |
| 1 | $0.38 \pm 0.02$ | $0.12 \pm 0.01$ | $0.26 \pm 0.03$ | $0.38 \pm 0.01$ |
| 2 | $0.59 \pm 0.05$ | $0.24 \pm 0.02$ | $0.40 \pm 0.03$ | $0.50 \pm 0.02$ |
| 3 | $0.71 \pm 0.05$ | $0.32 \pm 0.02$ | $0.49 \pm 0.02$ | $0.55 \pm 0.02$ |
| 4 | $0.74 \pm 0.04$ | $0.37 \pm 0.02$ | $0.52 \pm 0.01$ | $0.55 \pm 0.02$ |
| 5 | $0.72 \pm 0.04$ | $0.40 \pm 0.02$ | | |
| 6 | $0.69 \pm 0.05$ | | | |
| 7 | $0.66 \pm 0.05$ | | | |
| 8 | $0.66 \pm 0.07$ | | | |

CsCl has a high solubility in water, up to 7-7.5 M (40). However, we observe a sudden but reversible collapse of the DNA in 6 M CsCl solution when lowering the force down to 0.5 pN (**Supplementary Figure S7**). Therefore, we limited our measurements to concentrations of ≤ 4 M CsCl, for which no DNA collapse was observed, and recorded rotation curves at *F =* 0.5 pN, instead of 0.25 pN used for the other ions (**Supplementary Figure S6**). In this range, we observed changes in twist consistent with previous observations: At low concentrations (100 mM), Cs^+^ induces the largest twist of all monovalent ions tested, even larger than Li^+^. The twist induced by Cs^+^ is surpassed by Li^+^ at intermediate concentrations but continues to exceed both NaCl and KCl. At high concentrations (> 1 M), the twist induced by Cs^+^ continues to increase and appears to plateau around 4 M, similar to the trend observed for Li^+^. Overall, the change in twist is well described by a square root dependence up to 2 M for all salts studied, in good agreement with previous measurements (**Supplementary Figure S8**). However, for concentrations ≥ 4 M we observe clear deviations from this model and the observed change in twist clearly falls below the predictions of the square root dependence (**Supplementary Figure S8**). We note that for all tested conditions, the increase in twist compared to approximately physiological conditions is ≤ 0.8 degree/bp, which is well below the twist suggested for C-form DNA.

### Changes of the slope in the plectonemic regime

The DNA tether extension decreases linearly with increasing (absolute) linking number in the plectonemic regime and the slope in this regime provides a measure for the size of the plectonemes (71,72). We observe changes in the slope of the decrease in extension with applied turns (**Figure 1B and Supplementary Figure 10**), in good agreement with previous measurements in magnetic tweezers (73,74). At low salt concentrations (≤ 3 M), the slopes are equal in magnitude for positive and negative plectonemes for all salts investigated (**Figure 1B** and **Supplementary Figure S9**). Interestingly, at higher salt concentrations, the observed slopes remain mostly within experimental error but tend to be larger for positive than for negative supercoils, indicating possible -overall small-changes in helix or plectoneme geometry. We note that these changes are unlikely to be due to (partial) denaturation of the helix, since the curves were recorded at very low forces (*F =* 0.25 pN; except for CsCl), where torque-induced melting is unlikely to occur, in particular since the pH of the solution remains close to neutral (**Supplementary Figure S10**).

### Circular dichroism spectra suggest changes in DNA conformation at high salt concentrations

To further study the changes in DNA induced by high salt concentrations and in particular to compare to previous reports (30,31,33,36), we measured circular dichroism (CD) spectra. For 100 mM monovalent alkaline ion concentrations the CD spectra present three bands above 210 nm, with positive bands around 225 nm and 280 nm and a negative peak around 250 nm (**Figure 2A and Supplementary Figure S11),** in line with the well-established CD spectra of canonical B-form DNA (75). At 100 mM salt concentration, the band at 280 nm band exhibits some dependence on ion identity (**Supplementary Figure S11**), in line with previous observations (33).

**Figure 2.**
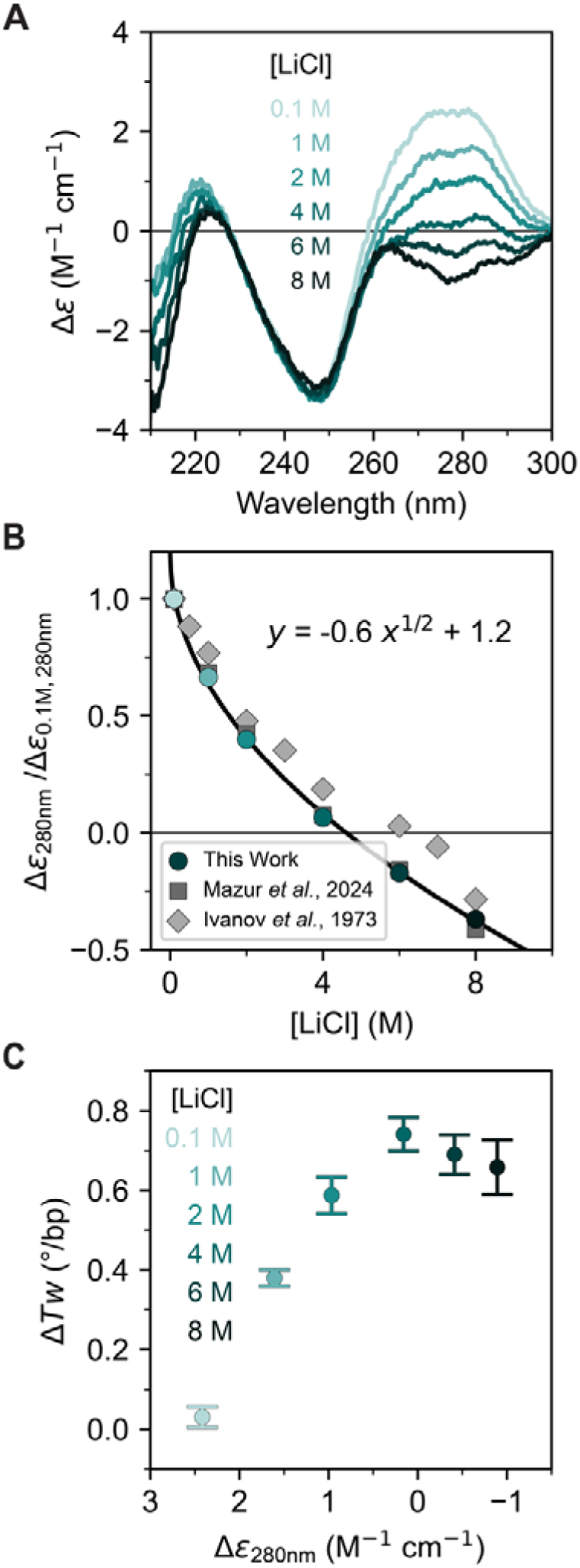
Effect of increasing LiCl concentration on CD spectra of DNA. A) Circular dichroism spectra of DNA (λ-DNA, 40 μg/mL) for increasing LiCl concentrations (from light to dark teal). Spectral changes are observed around the 280 nm band as the LiCl concentration is increased. B) Differences in the molar absorption at 280 nm for increasing LiCl concentrations. Data are normalized to the 100 mM LiCl condition. The solid line is a fit of a square root concentration dependence. Circles are from the data in panel A. Squares and diamonds are data taken from Refs. (36) and (33), respectively. C) Comparison of the change in DNA twist (Figure 1G) and the difference in the molar absorption at 280 nm (panels AB) for increasing LiCl concentrations. For additional data see **Supplementary Figure S11**. Symbols are the mean ± SD from at least 11 molecules for Δ*Tw* compared to the difference in molar absorbance values at 280 nm from the spectra in panel A.

In the presence of high concentrations of LiCl (1-8 M), the band around 280 nm is systematically reduced and a sign change occurs above 4 M LiCl. The effect of salt on this spectral region of DNA has been widely reported and our experimental results are in good agreement with the observed changes in CD in previous work, using different DNA constructs with different sequences (30,31,33,36) (**Figure 2B**). Control measurements at different DNA concentrations show no significant dependence on DNA concentration (**Supplementary Figure S11**), suggesting that intermolecular interactions, finite concentration effects, or aggregation are unlikely to play an important role in the changes in the CD spectra for the linear DNA used in our study. The change in the CD band at 280 nm follows a power law and scales with the square root of the salt concentrations over the entire investigated salt range, in excellent agreement with the observed changes in CD in recent work on linearized plasmids (36) (**Figure 2B**).

To characterize the changes in DNA structure revealed by CD spectroscopy, we analyzed the data by singular value decomposition (SVD) (see **Supplementary Figure S12** for details). We find that the CD spectra are well described by two components with large singular values, with a third somewhat marginal signal-containing component. Unlike protein or RNA folding (76–78), the weights of our basis functions show a gradual evolution, lacking a clear sigmoidal transition. Interestingly, the coefficient associated with the second basis function displays a power law dependence similar to the CD intensities observed at 280 nm. Overall, the SVD analysis indicates gradual changes in the CD spectra and not the presence of two characteristic spectra or states.

The trend in the CD spectra, in particular at 280 nm, is similar to the scaling of DNA twist observed for alkaline salts up to 1 M (5), but unlike the behavior of DNA twist as determined by MT measurements at high salt concentrations (**Figure 1**). From the MT measurements, we find the maximum change in twist for all four salts studied, LiCl, NaCl, KCl, and CsCl to be at 4-5 M (**Figure 1G**, **Table 1**). For LiCl, we directly compare the change in twist with the change in CD at 280 nm (**Figure 2C**). Below 4 M a similar trend in both Δε_280nm_ and Δ*Tw* is observed, as is for other salts when data from previous work is compared (**Supplementary Figure S13**) suggesting a correlation between helical twist and CD intensity at 280 nm. However, above 4 M the decrease in Δε_280nm_ is no longer paired with an increase in DNA twist, as it remains at ≈0.7 degree/bp for LiCl. Together, these observations suggest that further changes in the CD spectra, including the sign change, beyond 4 M are not associated with a continued increase in overall DNA twist and result from different types of molecular changes.

### MD simulations predict the change in DNA twist for high LiCl concentrations in qualitative agreement with experiments

We performed unbiased all-atom MD simulations using a 33 bp DNA duplex in solution with increasing LiCl concentrations up to 7.8 M (10 m on the molality scale). The DNA constructs used in the MD simulations are necessarily much shorter than the DNA used for MT measurements, since all-atom MD simulations with explicit water and ions of ≥ kbp DNA are currently beyond the scope of computational capabilities. However, previous work has shown that DNA twist can be reliable determined from simulations of DNA oligomers as used here and systematically compared to MT measurements on much longer DNA or RNA constructs (5,70,79,80). In addition, DNA conformations can be strongly influenced by sequence (81–83). In our previous work (5), we demonstrated that sequence-dependent effects are evident in oligonucleotides up to 15 bp in length, but these effects average out in longer constructs, such as the 33 bp duplex analyzed here.

We chose Li^+^ as cation, as it shows the largest DNA twist in the experiments. We compared several DNA force fields (parmbsc1, OL21, TUMUC1, and CHARMM36) and the solvent representation by Strelnikov et al. for which B-to-C transitions were observed for poly(AC) sequences (28).

The predicted change in DNA twist induced by high LiCl concentrations relative to 0.10 M KCl varies markedly across different force fields (**Supplementary Figure S14**). Notably, TUMUC1 and CHARMM36 predict a decrease in twist relative to 0.10 M KCl, contrary to experimental observations. Still, all force fields, except CHARMM36, capture the experimentally observed increase in twist with rising LiCl concentration relative to 1 M LiCl (**Figure 3A**). The deviations from experiments with CHARMM36 are linked to frequent openings of the terminal base pairs observed for all salt concentrations (**Supplementary Figure S15**). For TUMUC1, the discrepancy arises from excessive Li□ binding to phosphate oxygens and nucleobase donor atoms (**Supplementary Figure S15**). The deviations with TUMUC1 are not inherent shortcomings of the force fields themselves but reflect an unfavorable combination of DNA and ion force fields. Still, neither DNA force field is recommended for simulations of DNA in high salt conditions.

**Figure 3.**
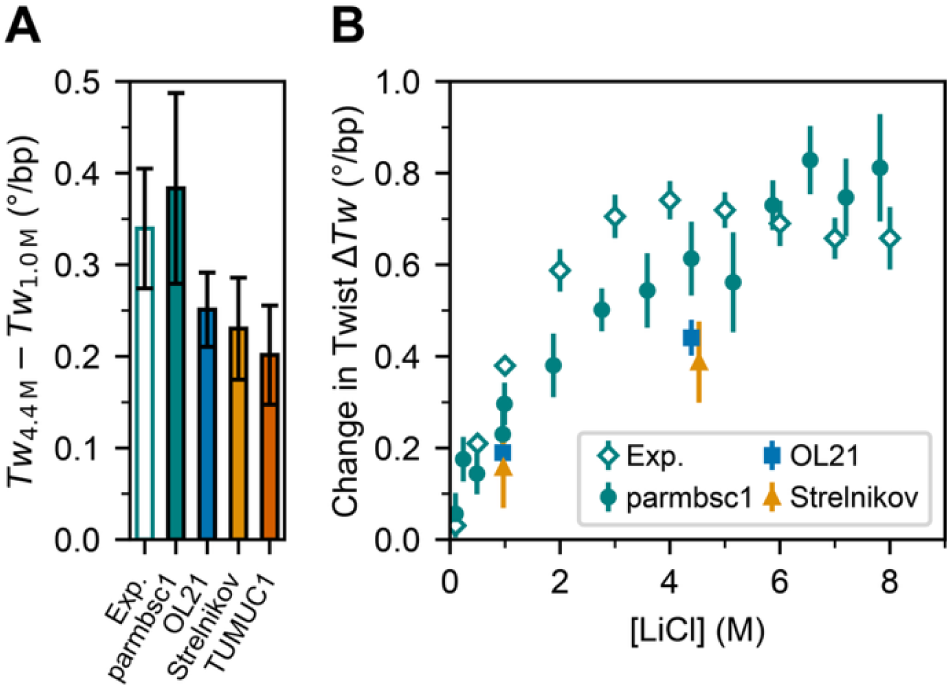
Change in twist from MD simulations for different force fields and in dependence of the LiCl concentration. A) Increase in twist between 1 and 4.4 M LiCl from experiments and different force fields: parmbsc1(39), OL21 (38), parmbsc1 with Strelnikov solvent representation (28) and TUMUC1 (29). B) Change in twist as a function of LiCl concentration relative to 100 mM KCl from experiments and selected force field simulations. Experimental data points are from Figure 1G and data points from simulations correspond to the mean ± SEM from six independent simulation runs.

Among the tested force fields, parmbsc1 combined with Mamatkulov–Schwierz ion parameters shows the best agreement with experimental data (**Figure 3** and **Supplementary Figure S14**) and is therefore used for the subsequent analysis. It correctly reproduces both the sign and approximate magnitude of twist increase up to ≈0.8 °/bp compared to 0.10 M KCl and provides qualitative agreement with the observed concentration dependence. The agreement between simulations and experiments improves further when differences in ion activities are accounted for by using activity derivatives, see below.

### C-form DNA is unstable in high LiCl concentrations and rapidly converges to B-form in MD simulations

We initiated half of our simulations in an ideal B-form (low twist) and the other half in the C-form (high twist). In all cases, i.e., for all force fields and solvent representations, the C-form is not stable, and the twist decreases rapidly to the lower values characteristic of B-form DNA in the course of the simulations. Specifically, after equilibration, the twist is at least around 2.5 °/bp lower compared to an ideal C-form structure, independent of starting configuration. Even at the highest 7.8 M concentration (parmbsc1/Mamatkulov-Schwierz), the twist for the C-form quickly decayed within a few nanoseconds to a value typical for B-DNA (**Figure 4A**).

**Figure 4.**
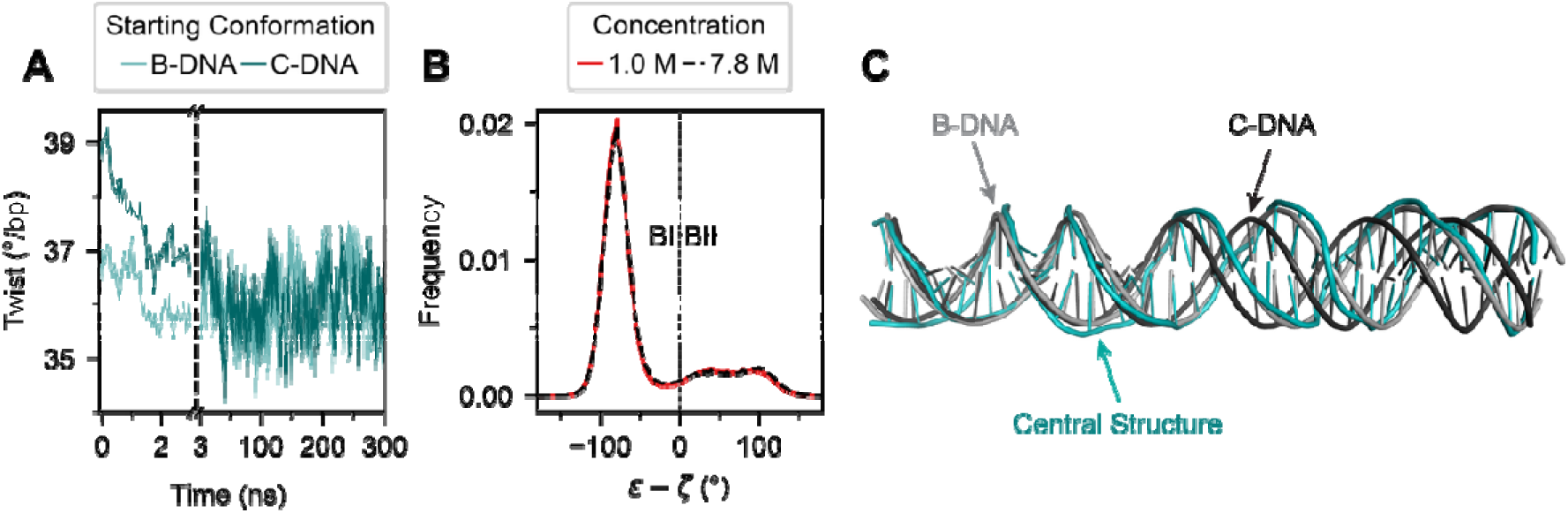
C-DNA is unstable in high LiCl concentrations and converges to B-DNA in MD simulations. A) Time series of twist in 7.8 M LiCl initiated from an ideal B- and C-DNA structure. The twist for the C-form quickly decays to the value of B-DNA. B) Histograms of the dihedral angles difference ε-ζ for 1 M and 7.8 M concentrations. The dashed vertical line indicates the separation between BI (ε-ζ < 0°) and BII (ε-ζ ≥ 0°) populations. C) Central structure of the dominant cluster (90% of structures) from 7.8 M concentration simulations (teal) superimposed with ideal models of B-(light gray) and C-form DNA (black). The structures were aligned using the first 10 bps on the left. All simulations were performed with parmbsc1/Mamatkulov-Schwierz force fields.

BI/BII populations provide an alternative means to distinguish between B- and C-form DNA, as the C-form of DNA has been experimentally associated with elevated BII populations at high salt concentrations (28,84). To further confirm that C-DNA does not form in our simulations, we analyzed these populations in detail for the parmbsc1/Mamatkulov-Schwierz simulations. The distributions of the ε–ζ dihedral angle difference, where negative values indicate BI and positive values BII populations, are nearly identical at 1 M and 7.8 M LiCl, with no systematic shift toward BII at higher concentrations (**Figure 4B**). Consistently, the BII population remains within 21–23% across all conditions as is typical for B-DNA and far below the > 40% expected for C-DNA (28).

In addition, we clustered structures from simulations at 7.8 M LiCl that started in C-DNA and compared the central structure to idealized B- and C-forms (**Figure 4C**). The central structure of the largest cluster (90% of all structures) aligns more closely with the ideal B-form (RMSD 2.8 Å) than with the C-form (RMSD 5.5 Å) further supporting the absence of C-DNA in the simulations. In summary, twist, BI/BII populations, and structural alignment show consistently that C-form DNA is not stable in the simulations for different force fields, even at LiCl concentrations up to 7.8 M, and rapidly converges to B-form.

### MD simulations reproduce the experimental change in DNA twist as a function of the activity derivative

The change in twist obtained from simulations as a function of salt concentration shows two notable deviations from the experimentally determined values: First, the initial increase in twist in the simulations is less steep compared to experimental results (**Figure 3B**). Second, while the experimental values saturate between 4 and 5 M salt concentration, the twist from simulations continues to increase and saturates at about 6.5 M. The observed discrepancies suggest that while MD simulations capture the sign and magnitude of DNA twist changes with high LiCl concentrations, they fail to describe the concentrations dependence quantitatively. We next tested whether deviations between ion activities from simulations and experiments can explain the apparent deviations in concentration dependence.

Activity coefficients and their derivative provide direct insight into ion–ion and ion–water interactions (51,85). In simulations, activity derivatives can be directly accessed via Kirkwood–Buff theory (53,54), and we used this approach to compare the activity derivatives from our simulations to experimental data (**Supplementary Methods** and **Supplementary Figures S16 and S17**). The comparison between the experimental and calculated activity derivative as function of the salt concentration (**Figure 5A**) shows that the activity derivative in the simulations is too low for concentrations larger than 2.5 M. An underestimation of the activity derivative indicates that ion pairing between Li^+^ and Cl^-^ is overestimated in the simulations (17,51).

**Figure 5.**
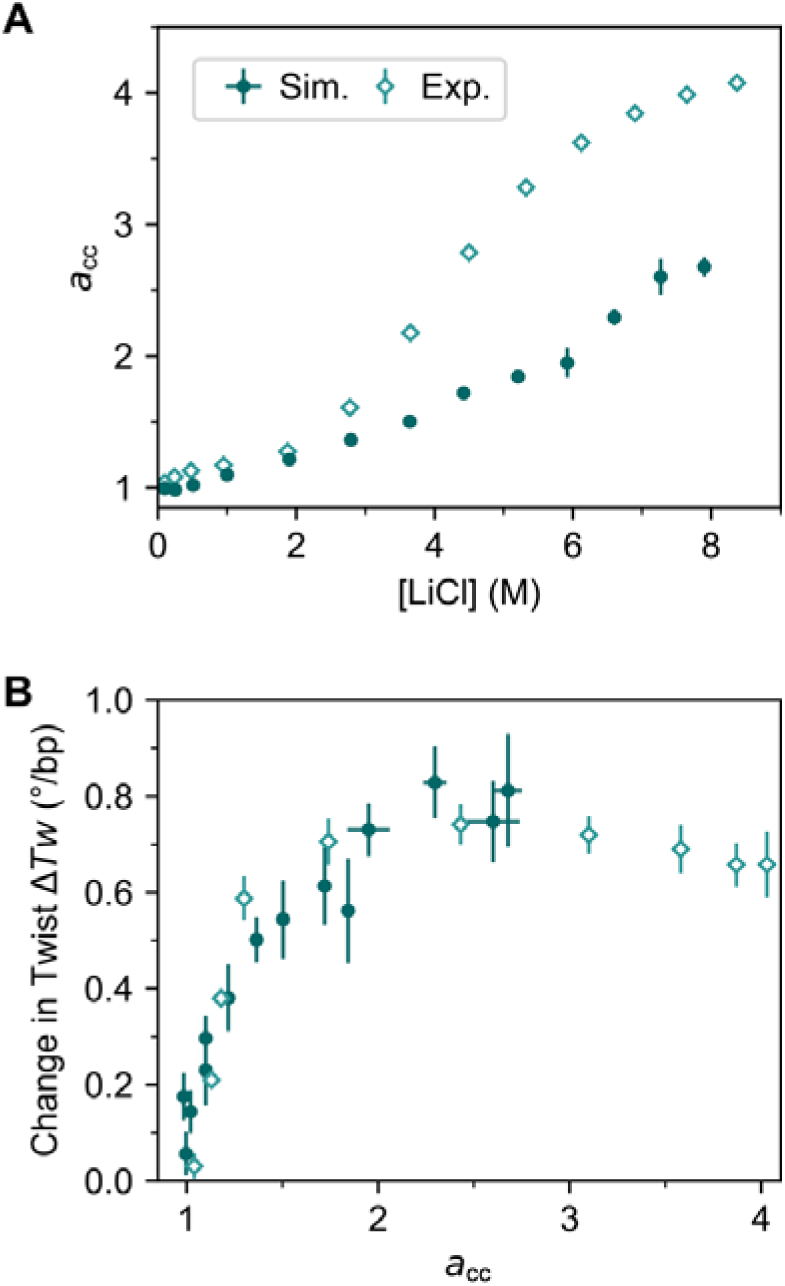
Change in twist as function of the activity derivative. A) Activity derivative a_CC_ as a function of the LiCl concentration from experiments (102) and current simulations using Kirkwood-Buff theory. B) Change in twist as a function of the activity derivative a_CC_ obtained from the a_CC_ -concentration relation shown in A) from experiments and simulations with parmbsc1 combined with Mamatkulov–Schwierz ion force field, and the TIP3P water model. Simulation points and error bars for a_CC_ indicate the mean and SEM from three independent simulations.

To account for the deviations, we compare experiments and simulations under conditions where anion-cation interactions are matched, using activity derivatives as the basis for comparison. Using activity derivates instead of concentrations, we find significant improvement and quantitative agreement between experiments and simulations (**Figure 5B**). Notably, the simulations capture the experimentally observed non-linear increase of twist including the steep increase at low activity derivatives, the slowing of twist change in the intermediate range and the saturation at activity derivatives larger than 2 (corresponding to an experimental concentration of around 4 M).

We conclude that accurate predictions of DNA twist at high salt concentrations require proper treatment of both ion–ion and ion-water interactions as well as ion binding while an imbalance leads to deviations between simulations and experiments. We find that comparing DNA properties as a function of activity derivative rather than concentration provides a more reliable and accurate approach. The observed deviations suggest, further, that refining the Li□–Cl□ interactions beyond standard combination rules at high salt concentrations, analogous to previous efforts under low-salt conditions (85–87), likely represents an important next step for accurate high-salt MD simulations.

### Mechanistic model for DNA twist in the high salt regime

To provide molecular insight into the non-linear increase and saturation of DNA twist with increasing activity derivative, we analyzed characteristic structural properties of DNA (**Figure 6**). Overall, we find that changes in twist correlate strongly with the DNA radius, sugar pucker angle, and helical rise (**Supplementary Table S5**), consistent with previous findings in the low-salt regime (5,6). In particular, the trends observed at low salt concentrations persist into the high-salt regime: as salt concentration increases, more Li□ ions adsorb to the DNA backbone (**Figure 6F,G**) leading to enhanced electrostatic screening. This screening reduces phosphate–phosphate repulsion and results in a decrease in DNA radius (**Figure 6A**). The narrowing of the helix is compensated by an increase in the sugar pucker angle (**Figure 6B**) and a decrease in helical rise (**Figure 6C**). Notably, the change in twist with increasing activity derivative are non-linear: The initial sharp increase is followed by a slowing and eventual saturation at very high concentrations, for activity derivative ≥ 2, corresponding to ≈4 M salt (**Figure 5B**). The same trend is also observed in the changes in radius and sugar pucker angle (**Figure 6A,B**).

**Figure 6.**
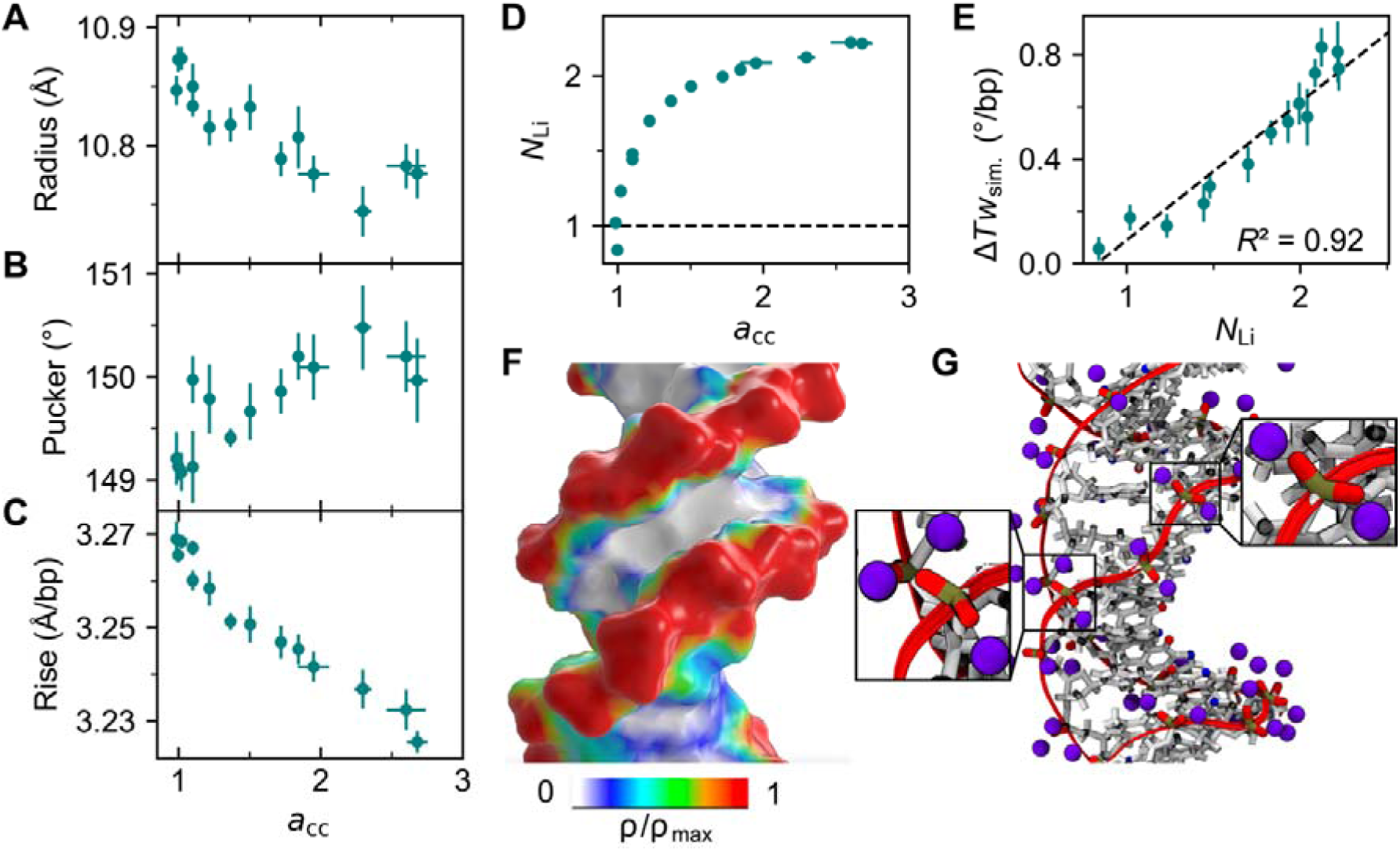
Structural properties of DNA and ion adsorption in the low and high salt regime. Change of structural properties of DNA with increasing activity derivative *a*_cc_: A) DNA radius, B) sugar pucker angle, C) helical rise. D) Number of inner-sphere Li^+^-ions per phosphate group N_Li_ as a function of the activity derivative a_CC_ using the a_CC_ -concentration relation from Figure 5A. The dashed line indicates neutralization of the negative phosphate charge. E) Change in twist from simulations LTw_Siffi_ as a function of N_Li_. F) Three-dimensional ion distributions projected on the DNA surface. (G) Simulation snapshot illustrating the Li^+^ ions (purple) within 3 Å of the non-bridging oxygen atoms of DNA from simulations at 4.4 M LiCl concentration. Zoom-in on selected Li^+^ binding sites: Two Li^+^ in inner-sphere configuration with the phosphate oxygens in the upper right and three Li^+^ ions per phosphate group in the lower left. All simulations were done with the parmbsc1/ Mamatkulov–Schwierz force fields. Data points in panels A-E correspond to the mean ± SEM from six independent simulation runs, except for the activity derivative, where the data are from three independent simulations.

The underlying mechanism is closely linked to the number of specifically adsorbed Li^+^ ions (**Figure 6D** and **Supplementary Figure S18**). In the low salt regime, the number of adsorbed cations increases rapidly, approaching full compensation of the negative DNA charge near activity derivative *a*_cc_ = 1 (corresponding to ≈1 M) with one Li^+^ per phosphate group (dashed line in **Figure**□**6D**). Beyond this point, charge reversal occurs, and the local effective DNA charge becomes positive, indicating overcharging consistent with previous observations for other monovalent cation-DNA interactions (88). Overall, ion binding slows as the system approaches a stoichiometry of two Li^+^ ions per phosphate group. Here, on average two Li^+^ ions are bound per phosphate group, each forming an inner-sphere complex with the phosphate oxygens (**Figure**□**6G**). For two Li^+^ ions per phosphate group (N = 2) the DNA charge is exactly reversed. Above this point, the number of adsorbed ions saturates at about N_Li_ = 2.1. Further ion accumulation is hindered by electrostatic repulsion, which limits the proximity of additional Li^+^ ions. Nevertheless, a third Li^+^ ion can still bind (**Figure 6G**), leading to a saturation level slightly above two ions per phosphate (**Figure 6D**).

Finally, the change in twist correlates strongly with the number of specifically adsorbed Li^+^ ions (**Figure 6E and Supplementary Table S5**). This suggests that in the low salt regime, increased electrostatic screening increases twist, sugar pucker and decrease radius and rise. In the intermediate salt regime, ion binding slows as the system reaches two Li^+^ ions per phosphate group and hence the changes in twist and helical properties slow down accordingly. Finally, the onset of the ion adsorption plateau coincides with the saturation of twist, suggesting that high-affinity Li^+^ binding is limited electrostatic repulsion between the cations, which in turn defines an upper bound for DNA twist.

## CONCLUSIONS

Our results demonstrate that the increase in DNA twist with increasing salt concentration, previously documented at lower concentrations, persists well beyond 1 M for all tested monovalent ions, LiCl, NaCl, KCl, and CsCl. This increases in DNA twist, as evidenced by shifts in magnetic tweezers rotation curves, reaches saturation between 4-5 M (**Figure 1G**). Notably, the ion-specific differences in DNA twist observed at lower concentrations persist even at the highest salt levels tested and the twist values at saturation follow the same trend: Li^+^ > Cs^+^ > K^+^ > Na^+^.

To complement the magnetic tweezers measurements, we performed all-atom MD simulations, focusing on LiCl, which produced the largest experimental change in DNA twist. Using selected force fields (parmbsc1 for DNA, the Mamatkulov–Schwierz model for ions, and TIP3P for water) the simulations reproduce the twist increase up to high ion concentrations qualitatively. In contrast, other combinations of DNA and solvent force fields exhibit notable shortcomings in capturing the experimental trends.

While the simulations qualitatively reproduce the twist increase, they systematically underestimate the ion activity and its derivative above 2.8 M, suggesting that they overestimate the Li^+^-Cl^-^ interactions. We find that comparing simulations and experiments using activity derivatives rather than concentrations significantly improves the agreement, capturing both the nonlinear twist increase and its saturation (**Figure 5B**). This highlights the critical importance of accurately modeling ion-ion and ion-water interactions at high salt concentrations, where standard force fields are typically not optimized. For instance, an activity derivative of ≈ 2.7, corresponding to 4–5 M LiCl experimentally, reveals a two-fold discrepancy between simulated and experimental activity derivatives. Improving force field accuracy will require refinement of ion–ion, ion–water and ion-DNA interactions, specifically through optimizing the cation–anion combination rules at high salt based on experimental data (85–87).

Our results directly address the question of whether C-form DNA exists at high salt concentrations in free solution: Notably, we do not observe a shift of ≈2 °/bp twist increase relative to low salt that has been reported for C-form DNA in fiber diffraction studies. Instead, our measurements reveal a maximal twist increase of only ≈0.75 °/bp, which is consistent, within error, with previous ultracentrifugation measurements in 6.2 M LiCl (34). These experimental findings are fully corroborated by our MD simulations, which reproduce the observed increases of twist but likewise show no evidence for the C-form in solution even at high salt concentrations. Instead, simulations initiated in either canonical B-form or C-form rapidly converge to B-form, with twist values matching experimental data. While different DNA constructs were used for single-molecule, bulk, and computational experiments due to the specific requirements of each technique, the heterogeneous, “random” sequences used in our assays are considered representative of generic DNA sequences. Therefore, together, our findings suggest that in aqueous solution, even at very high monovalent salt concentrations, DNA with genomic or random sequences remains in the B-form and does not adopt the C-form.

Previously, continuous changes in CD spectra at increasing salt concentrations were interpreted as evidence for the presence of C-form DNA in solution. Our CD spectra are in excellent agreement with these earlier reports. Notably, changes in the band around 280 nm up to 4–5 M concentrations of monovalent alkaline ions, closely correlate with the increase in DNA twist observed directly by magnetic tweezers and reproduced in our MD simulations. Interestingly, however, both our data and previously published CD spectra show continued spectral shifts at 280 nm even beyond 4–5 M salt, a regime in which the DNA twist, as measured by magnetic tweezers, has already plateaued.

The fact that Δ*ε* at 280 nm undergoes further changes at salt concentrations where the twist has plateaued and remains constant implies that CD does not directly report on DNA helical twist. It is currently unclear what molecular features contribute to this detail of the CD spectra precisely. One possibility are the continued changes in the ion binding pattern including the increasing number of Li^+^ ions bound to the nucleobases (**Supplementary Figure S18**).

In this context, a promising direction for future research is the development of computational approaches capable of predicting CD spectra directly from MD-generated DNA structures. For proteins, approximate knowledge-based tools such as SESCA (89) or DiChroCalc (90), are already available. For DNA and other systems less well-characterized by CD, however, more rigorous first-principles methods such as time-dependent density functional theory or exciton coupling models are likely required (91,92). In addition, a complementary future direction would be to employ other -ideally high-resolution-structural probes of DNA conformations and dynamics, for example FRET (93–96), (anomalous) X-ray scattering interference (97–99), or NMR spectroscopy (100,101), to further experimentally probe DNA structure in solution at very high ionic strengths.

In summary, our data quantitatively describe how DNA twist responds to very high salt concentrations and provide a critical test of molecular force fields in this previously unexplored regime. They offer a robust experimental benchmark for validating simulations and establish a baseline for modeling DNA at extreme ionic conditions, with implications for understanding nucleic acid behavior in extremophiles and prebiotic environments.

## DATA AVAILABILITY

The data underlying this article are available freely in the repository Zenodo at https://doi.org/10.5281/zenodo.17093173

## ACKNOWLEDGEMENTS

We thank Alexey Mazur for alerting us to the question of C-DNA formation at high LiCl concentrations and for useful discussions, Dave van den Heuvel, Elleke van Harten for laboratory support, Eefjan Breuking and Christian Kaiser for support with CD measurements, and Martin Zacharias for fruitful discussions.

