## Supplementary Information for "DNA Twist at High Alkaline Ion Concentrations: Evidence against C-Form DNA in Solution"

<u>**Content**</u>

**Supplementary Methods**

**Supplementary Tables S1-S5**

**Supplementary Figures S1-S18**

### Supplementary Methods

#### Determination of Activity Derivative from Simulations and Experiments

The activity coefficient $a_c$ is a factor to take the experimentally measured deviation of a mixture from ideal behavior into account. Similarly, the activity derivative $a_{cc}$ provides information on the balance between ion-ion and ion-water interactions (1,2). Kirkwood-Buff (KB) theory (3) was employed to compute this $a_{cc}$ of LiCl at various concentrations. Based on radial distribution functions between two species $i$ and $j$, the KB integrals $G_{ij}$ allow to determine a variety of thermodynamic properties (4):

$$G_{ij} = 4\pi \int_0^{\infty} \left[g_{ij}^{\mu VT}(r_{ij}) - 1\right] r_{ij}^2 \, dr_{ij}, \tag{1}$$

with $g_{ij}^{\mu VT}(r_{ij})$ being the radial distribution function of the two species $i$ and $j$ in the grand canonical ensemble and $r_{ij}$ being the center of mass distance between the two. Formally, the KB integrals are defined for infinite space and open systems (μVT), but MD simulations are typically realized for closed systems (NVT, NpT). Furthermore, due to the finite size of simulation boxes, also the integration cannot be carried out over all space. Therefore, the KB integrals are truncated and rescaled resulting in:

$$G_{ij} \approx 4\pi \int_0^{R} \left[g_{ij}^{NpT}(r'_{ij}\rho) - 1\right] r_{ij}^2 \, dr_{ij}, \tag{2}$$

in which the radial distribution function is rescaled via $g_{ij}^{NpT}(r'_{ij}\rho) = f(\rho) g_{ij}^{sim}(r_{ij})$ with a prefactor $f(\rho)$ that is adjusted to ensure the correct asymptotic behavior at large distances (5).

The activity derivative $a_{cc}$ is defined as:

$$a_{cc} = \left(\frac{\partial \ln a_c}{\partial \ln\rho_c}\right)_{p,T} = 1 + \left(\frac{\partial \ln \gamma_c}{\partial \ln\rho_c}\right)_{p,T} = \frac{1}{1 + \rho_c (G_{cc} - G_{co})}, \tag{3}$$

with the activity $a_c = \rho_c \gamma_c$, the cosolvent molar activity coefficient $\gamma_c$, and number density $\rho_c$.

For monovalent cations and anions, the required expressions are (2):

$$G_{cc} = \frac{1}{4}[G_{++} + G_{--} + 2G_{+-}] \text{ and } G_{co} = G_{oc} = \frac{1}{2}(G_{+o} + G_{-o}),$$

where +, -, and *o* denote cation, anion, and water oxygen, respectively.

For the simulations of LiCl with Mamatkulov-Schwierz parameters (2) in water at various concentrations, we employed cubic boxes with TIP3P water molecules (6) and inserted the corresponding numbers of cations and anions by replacing water molecules (**Supplementary Table S4**). The number of ions required for a given concentration was determined based on the number of water molecules in the simulation box. The equilibration protocol consisted of energy minimization followed by NVT and NpT equilibration. Energy minimization with the steepest descent algorithm used a maximum of 50000 steps. Subsequently, we used 1 ns of NVT and 1 ns of NpT simulations for further equilibration. In these simulations, the temperature of 300 K was maintained using the Berendsen thermostat with a coupling constant of 0.1 ps. In the NpT simulations, the isotropic Berendsen barostat with a coupling constant of 1 ps was employed to ensure a pressure of 1 atm.

For production runs, we used again NpT simulations at 300 K and 1 atm, but now by employing the velocity rescaling thermostat with a coupling constant of 0.1 ps and the isotropic Parrinello-Rahman barostat with a coupling constant of 5 ps. Three independent production simulations were carried out for 150 ns each. For determining the radial distribution functions, the first 5 ns of each production was discarded for equilibration. Furthermore, the radial distribution functions between ions were normalized in such a way that their average at large distances is 1. The KB integrals were evaluated numerically by employing the trapezoidal rule and with these integrals the activity derivative was determined via the final expression in Eq. (3). The three values for $a_{cc}$ per concentration were then used to evaluate the average and the standard error of the mean.

For obtaining experimental values of $a_{cc}$, we started with a selection of experimental activity coefficients taken from Hamer and Wu (7). First, the concentrations reported in molality were converted into molarity by employing a parabolic fit for molarity as a function of molality based on reference values (8) (**Supplementary Figure S17A**). To better approximate the required derivative, we performed a cubic fit of the activity coefficients as function of molarity in the concentration range from 0.09 to 8.40 M (**Supplementary Figure S17B**). We employed the equation of the cubic fit to generate closely spaced data points for $\ln \rho_c$ and $\ln a_c$ in the concentration range from 0.09 to 8.40 M (i.e. up to a molality of around 10 mol/kg) and used the central difference scheme for obtaining $a_{cc}$. These values for the concentrations of interest are shown in **Supplementary Figure S17C** and **Figure 5A**.

### Supplementary Tables

**Supplementary Table S1.** Fitting parameters for second-degree polynomial fits to the experimental index of refraction $n$ vs. salt concentration $x$ (in M) data (Supplementary Figure S2B).

| Model | Salt | Parameter | Value |
|---|---|---|---|
| $n=ax^2+bx+c$ | LiCl | $a$ | -0.0000922 |
| | | $b$ | 0.00866 |
| | | $c$ | 1.33296 |
| | NaCl | $a$ | -0.0002123 |
| | | $b$ | 0.00977 |
| | | $c$ | 1.33308 |
| | KCl | $a$ | -0.0002090 |
| | | $b$ | 0.00965 |
| | | $c$ | 1.33306 |
| | CsCl | $a$ | -0.0001123 |
| | | $b$ | 0.01247 |
| | | $c$ | 1.33318 |

**Supplementary Table S2.** Normalized difference in the molar absorption at 280 nm as a function of LiCl concentrations. Data are from the CD spectra in **Figure 2**.

| [LiCl] (M) | $\Delta\varepsilon_{280nm}$ |
|---|---|
| 0.1 | 2.42 |
| 1 | 1.61 |
| 2 | 0.97 |
| 4 | 0.16 |
| 6 | -0.41 |
| 8 | -0.89 |

**Supplementary Table S3.** Number of water molecules, cations and anions for the simulations of double-stranded DNA starting in B- or C-form, respectively, for the simulations with the parmbsc1 force field. Furthermore, the average volume from the production simulations and the resulting molarity are listed.

| B-Form | | | | | |
|---|---|---|---|---|---|
| Molality (m) | # $H_2O$ | # $Li^+$ | # $Cl^-$ | Volume ($nm^3$) | Molarity (M) |
| 1.00 | 66941 | 1271 | 1207 | 2089.5 | 0.96 |
| 2.00 | 64692 | 2396 | 2332 | 2061.0 | 1.88 |
| 3.00 | 62588 | 3448 | 3384 | 2037.6 | 2.76 |
| 4.00 | 60618 | 4433 | 4369 | 2018.3 | 3.59 |
| 5.01 | 58766 | 5359 | 5295 | 2002.3 | 4.39 |
| 6.01 | 57026 | 6229 | 6165 | 1988.9 | 5.15 |
| 7.01 | 55384 | 7050 | 6986 | 1977.5 | 5.87 |
| 8.01 | 53834 | 7825 | 7761 | 1967.6 | 6.55 |
| 9.01 | 52370 | 8557 | 8493 | 1958.8 | 7.20 |
| 10.01 | 50982 | 9251 | 9187 | 1950.8 | 7.82 |
| C-Form | | | | | |
| Molality (m) | # $H_2O$ | # $Li^+$ | # $Cl^-$ | Volume ($nm^3$) | Molarity (M) |
| 1.00 | 64997 | 1235 | 1171 | 2029.3 | 0.96 |
| 2.00 | 62811 | 2328 | 2264 | 2001.6 | 1.88 |
| 3.00 | 60769 | 3349 | 3285 | 1978.9 | 2.76 |
| 4.00 | 58855 | 4306 | 4242 | 1960.1 | 3.59 |
| 5.01 | 57057 | 5205 | 5141 | 1944.6 | 4.39 |
| 6.01 | 55367 | 6050 | 5986 | 1931.6 | 5.15 |
| 7.01 | 53773 | 6847 | 6783 | 1920.5 | 5.86 |
| 8.01 | 52269 | 7599 | 7535 | 1910.9 | 6.55 |
| 9.01 | 50847 | 8310 | 8246 | 1902.3 | 7.20 |
| 10.01 | 49501 | 8983 | 8919 | 1894.5 | 7.82 |

**Supplementary Table S4.** Number of water molecules, cations and anions for the simulations of ions in water (without DNA) to determine the activity derivatives. Furthermore, the average volume from the production simulations and the resulting molarity are listed.

| **LiCl in Water** | | | | | |
|---|---|---|---|---|---|
| Molality (m) | # $H_2O$ | # $Li^+$ | # $Cl^-$ | Volume ($nm^3$) | Molarity (M) |
| 0.10 | 2157 | 4 | 4 | 65.66 | 0.10 |
| 0.26 | 2145 | 10 | 10 | 65.47 | 0.25 |
| 0.52 | 2125 | 20 | 20 | 65.17 | 0.51 |
| 1.04 | 2087 | 39 | 39 | 64.64 | 1.00 |
| 2.01 | 2019 | 73 | 73 | 63.77 | 1.90 |
| 3.02 | 1953 | 106 | 106 | 63.03 | 2.79 |
| 4.02 | 1891 | 137 | 137 | 62.42 | 3.64 |
| 5.00 | 1835 | 165 | 165 | 61.94 | 4.42 |
| 6.03 | 1779 | 193 | 193 | 61.50 | 5.21 |
| 7.00 | 1729 | 218 | 218 | 61.15 | 5.92 |
| 8.00 | 1681 | 242 | 242 | 60.84 | 6.60 |
| 9.00 | 1635 | 265 | 265 | 60.56 | 7.27 |
| 10.02 | 1591 | 287 | 287 | 60.30 | 7.90 |

**Supplementary Table S5. Correlations of helical properties with helical twist, activity derivative ($a_{cc}$), and number of adsorbed ions (# Ions).** The helical properties consisted of the six base-pair step parameters with respect to a local helical frame (x-displacement, y-displacement, helical rise, inclination, tip, helical twist), major and minor groove widths and radius as defined in (9) as well as the average pucker and number of adsorbed ions (# Ions). Correlations are sorted in decreasing order of the absolute value of the correlation coefficient.

| **Helical Twist** $\Delta Tw$ | | $\mathbf{a_{cc}}$ | | **# Ions** | |
|---|---|---|---|---|---|
| Property | R value | Property | R value | Property | R value |
| # Ions | 0.960 | Helical Rise | -0.976 | Helical Twist | 0.960 |
| Major Groove | -0.958 | Helical Twist | 0.924 | Major Groove | -0.937 |
| Radius | -0.951 | Minor Groove | -0.870 | Helical Rise | -0.901 |
| Helical Rise | -0.947 | Radius | -0.864 | Radius | -0.877 |
| x-Displacement | 0.934 | # Ions | 0.859 | x-Displacement | 0.867 |
| Pucker | 0.847 | Major Groove | -0.858 | Pucker | 0.821 |
| Minor Groove | -0.777 | x-Displacement | 0.850 | Minor Groove | -0.697 |
| y-Displacement | 0.359 | Pucker | 0.779 | Inclination | 0.430 |
| Inclination | 0.353 | y-Displacement | 0.298 | y-Displacement | 0.406 |
| Tip | 0.146 | Inclination | 0.252 | Tip | 0.159 |
| | | Tip | 0.089 | | |

### Supplementary Figures

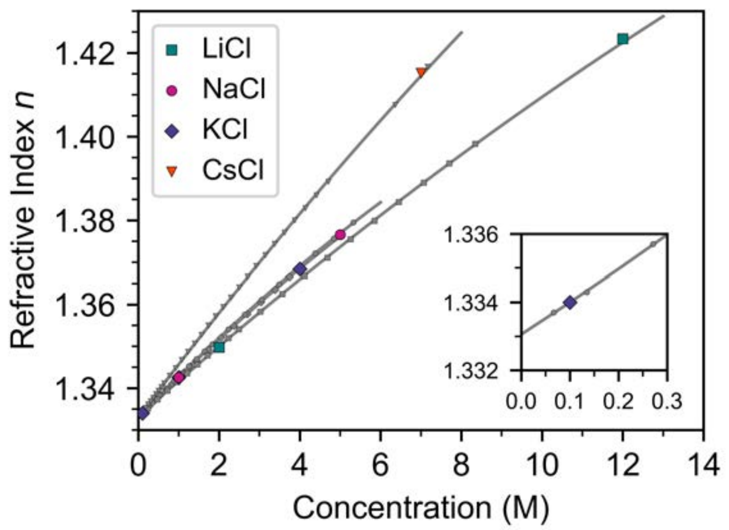


**Supplementary Figure S1. Concentration determination of salt stocks by index of refraction measurements.** We determined the index of refractions of several of our salt solutions using and Abbe refractometer (λ=589.3 nm, 21 ºC; Abbe Refractometer 3T, Atago). Measured data from this work are shown as colored symbols at the nominal concentration the stocks were prepared at by weighing the salts. Our measurements are in excellent agreement with the expected indices of refraction shown as grey symbols taken from (8). The solid lines are second-degree polynomial fit to the literature data.

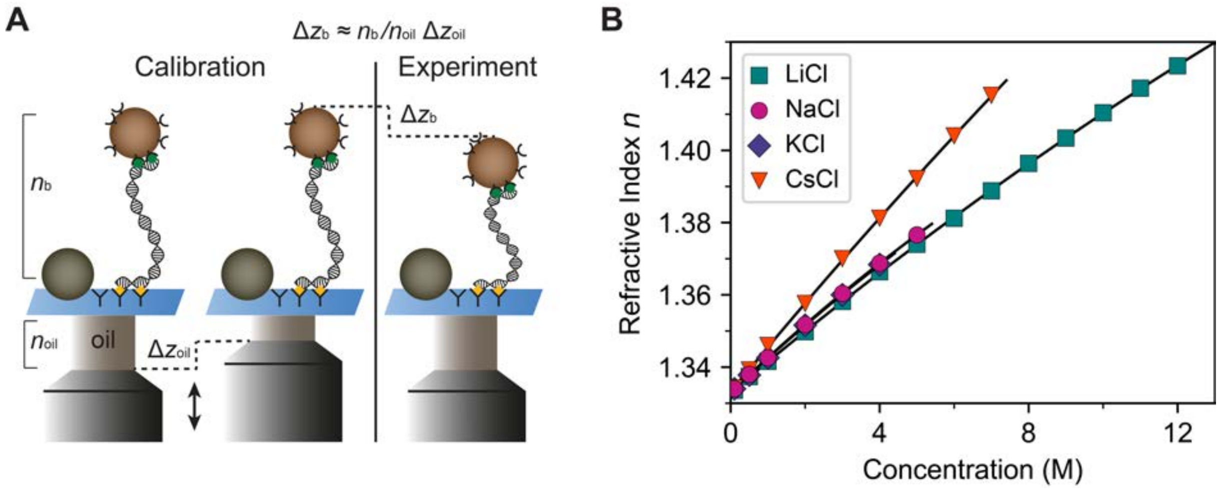


**Supplementary Figure S2. Effect of the index of refraction on MT measurements.** A) Schematic of the vertical displacement determination technique of magnetic tweezers. Prior to the experiment, calibration is performed where diffraction patterns are collected while vertically displacing the oil objective, i.e., a displacement of $\Delta z_{\mathrm{oil}}$. During measurements, the vertical position of the bead is correlated to the diffraction pattern. The absolute change in the vertical position depends on the ratio of the refractive index of the oil $n_{\mathrm{oil}}$ and of the buffer $n_{\mathrm{b}}$. We determine the absolute $\Delta z_{\mathrm{b}}$ according to $\Delta z_{\mathrm{b}} = n_{\mathrm{oil}} / n_{\mathrm{b}} \cdot \Delta z_{\mathrm{oil}}$. B) Refractive index of salt solutions in 10 mM Tris. The solid lines are second-degree polynomial fits to the data with coefficients shown in **Supplementary Table S1**.

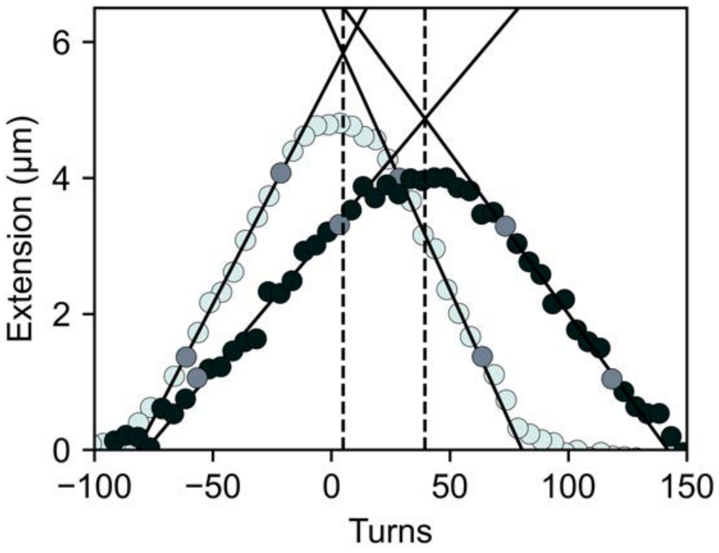


**Supplementary Figure S3. Determination of change in twist from MT rotation curves.** Extension vs. applied turns measurements for our 20.6 kbp DNA construct at an applied force of 0.25 pN for the 0.1 (light data points) and 7 M (black data points) LiCl conditions. In the plectonemic regime past the buckling points, the DNA tether shows a linear decrease in extension vs. turns upon addition of additional of positive or negative supercoils, respectively. The slopes in the buckling regimes at positive and negative turns are determined by fitting a linear relationship to the data in this range (grey circles indicate the beginning end of the fitting range), as described in Ref. (10). The centers of the rotation curves are determined as the intersection of the extrapolated fitted lines.

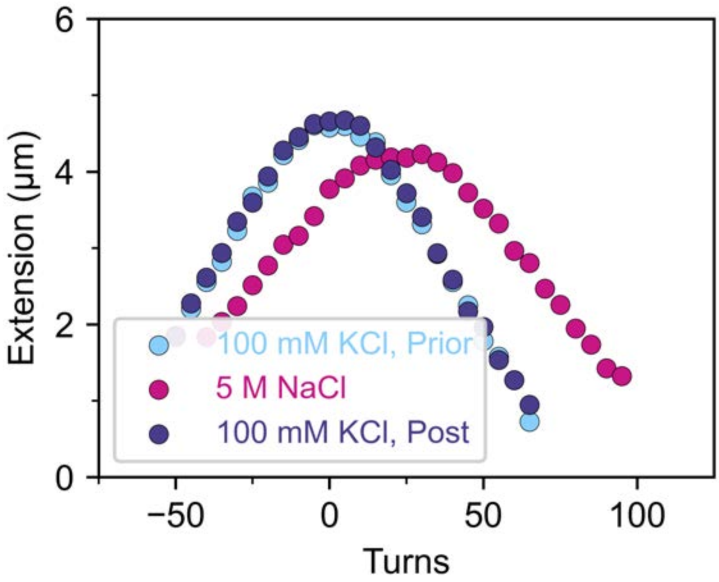


**Supplementary Figure S4. Reversibility of twist increase.** Rotation-extension curves collected in the 100 mM KCl reference condition and in 5 M NaCl. Upon return to the reference condition by exchanging 5 M NaCl for 100 mM KCl, the initial rotation curve is recovered within experimental error.

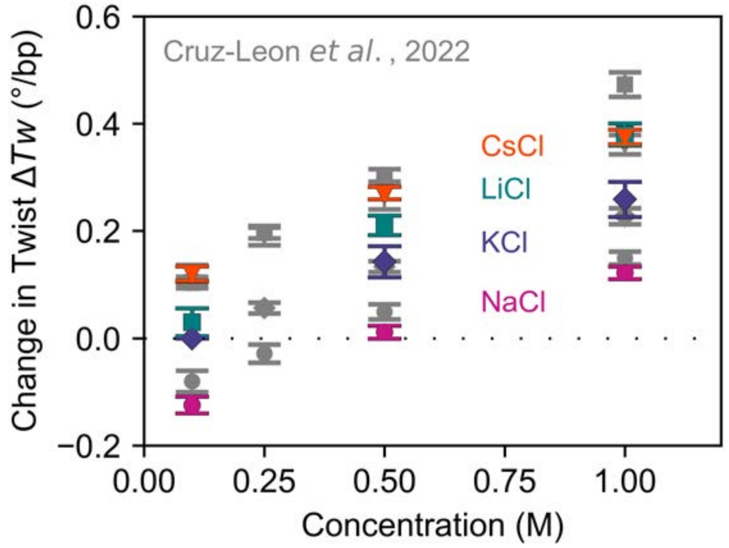


**Supplementary Figure S5. Comparison of MT twist measurements with Cruz-Leon *et al.*** Changes in DNA twist determined from MT measurements as described in Materials and Methods and Figure 1. Grey data are from Cruz-Leon *et al.* (11). Colored data points are from this work (same data as in Figure 1) and indicate the mean ± SD from at least 4 molecules. 100 mM KCl was used as a reference condition throughout. Overall, the data show good reproducibility. The data from this work and Cruz-Leon *et al.* for both CsCl and KCl are within experimental error. For LiCl and NaCl, we find a slightly, but systematically lower twist relative to 100 mM KCl in this work compared to the data of Cruz-Leon *et al.,* which could be due to a slightly mismatched reference condition. However, the observed deviations are small (< 0.05 deg/bp) and do not affect any of the conclusions drawn.

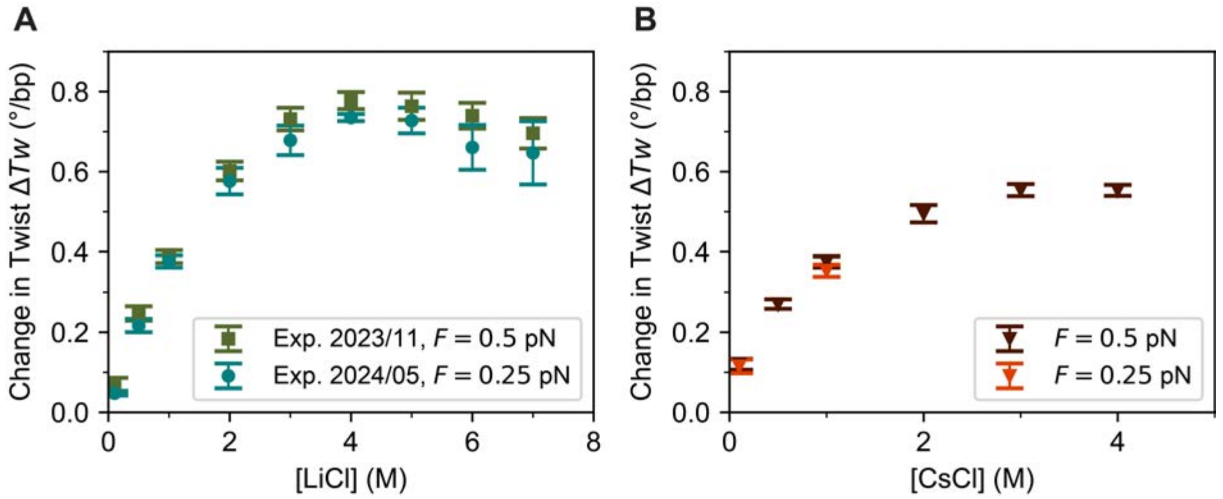


**Supplementary Figure S6. Reproducibility of MT measurements of DNA twist.** A) Changes in DNA twist determined from MT measurements as described in Materials and Methods and Figure 1. Comparison of a first LiCl data set, from a 10 M LiCl stock (≥99%, Sigma-Aldrich), recorded in 10 mM Tris pH 7.6 and using rotation curves at $F$ = 0.5 pN (2023/11), and a second data set, prepared as described in Materials and Methods, using rotation curves at $F$ = 0.25 pN (2024/05). The two data sets recorded for this work are in excellent agreement, within experimental error. Symbols are the mean ± SD of at least 2 molecules. B) Comparison of experimentally determined change in twist for CsCl at different forces. The change in twist determined from rotation-extension curves at $F$ = 0.25 pN are within error of values found for rotation-extension curves at $F$ = 0.5 pN for 0.1 and 1 M CsCl. Symbols are the mean ± SD of at least 10 molecules.

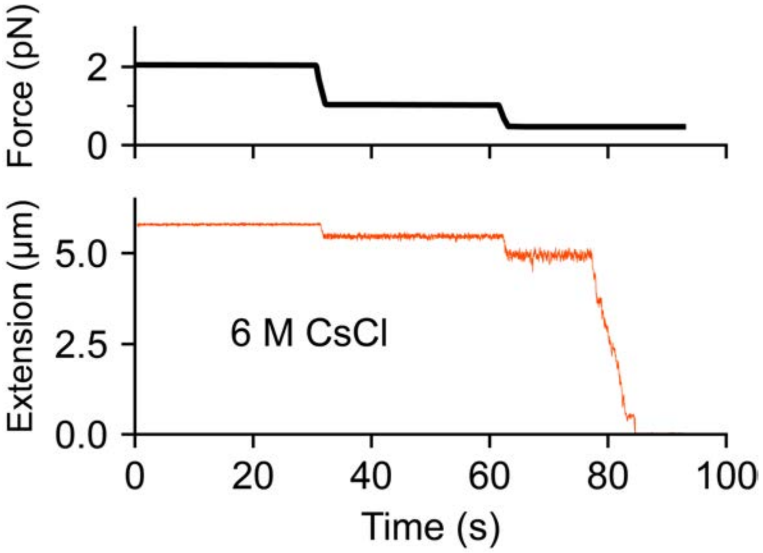


**Supplementary Figure S7. Collapse of DNA in the presence of 6 M CsCl.** Extension time trace of a 20.6 kbp DNA tether in 6 M CsCl. The applied force is indicated in the top panel. A collapse of the DNA tether is observed in the 6 M CsCl solution as the stretching force is lowered from 2 pN to 0.5 pN.

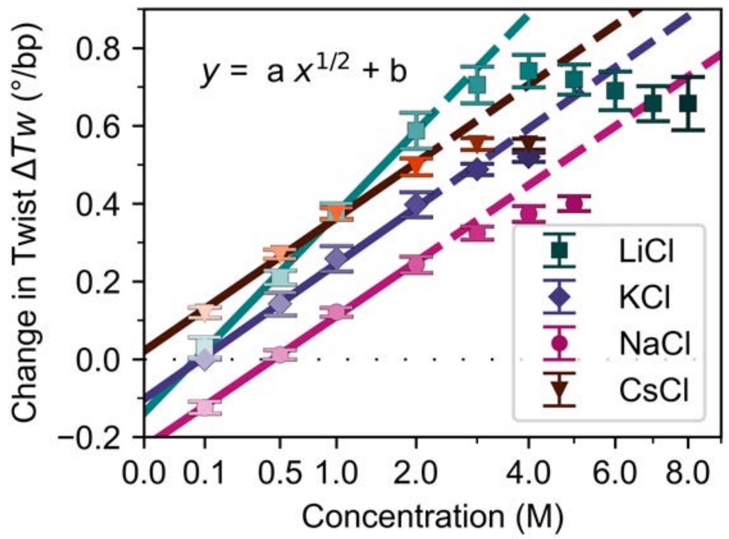


**Supplementary Figure S8. Changes in DNA twist with salt concentration as determined from MT measurements shown on a linear-logarithmic scale.** The symbols show the change in DNA twist determined experimentally (same data as Figure 1G). The solid lines indicate square-root dependencies fitted up to 2 M. Dashed lines are the extrapolation of the fits beyond a concentration of 2 M. The data are well described by the square root dependencies up to 2 M. At higher salt concentrations deviations from the square-root scaling are apparent and the experimental data fall below the square root trend lines.

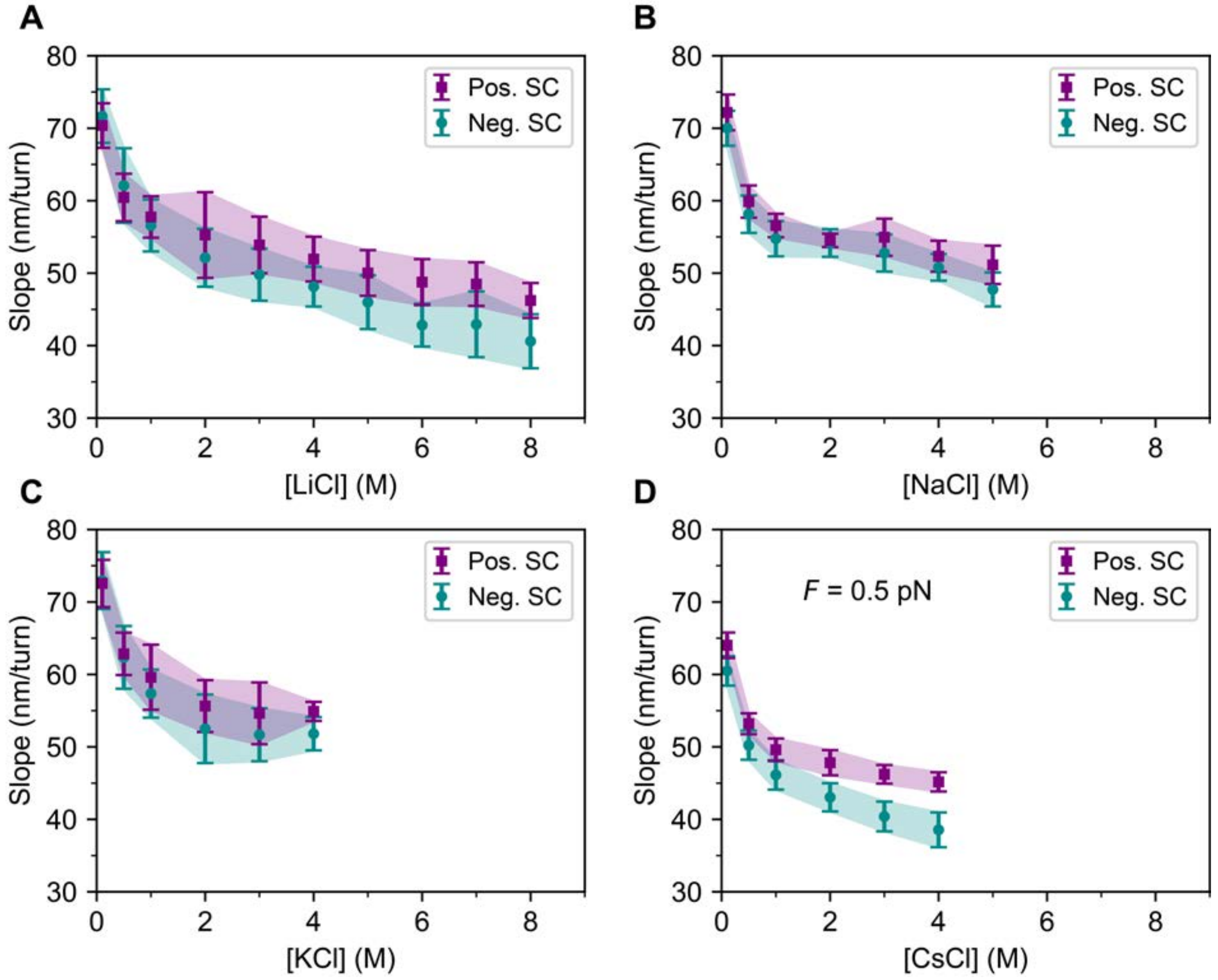


**Supplementary Figure S9. Fitted slopes of DNA extension vs. applied rotation in the presence of LiCl, NaCl, KCl or CsCl.** Extension vs. turns slopes in the plectonemic regime from fits to the rotation curve data for different concentrations of A) LiCl (same data as Figure 1B), B) NaCl or C) KCl, recorded at a force $F$ = 0.25 pN or D) CsCl, recorded at a force $F$ = 0.5 pN.

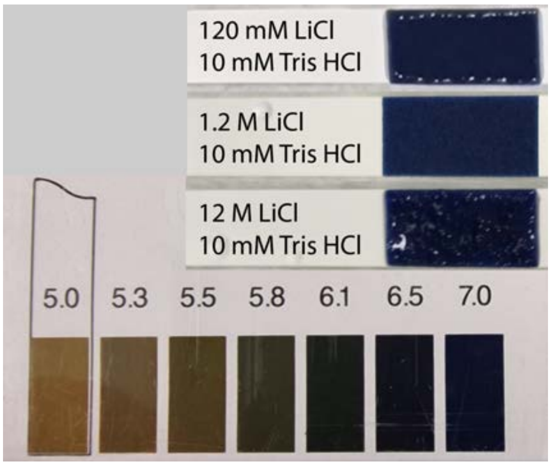


**Supplementary Figure S10. Measurements of solution pH at different LiCl concentrations.** Measurements of pH for a broad range of LiCl concentrations using pH indicator paper (Merck, Germany). Aqueous LiCl has been shown to remain close to neutrality but suggested to cause slight acidity at extreme conditions (12). The pH indicator strips show that solutions up to 12 M do not present noticeable acidity under our conditions.

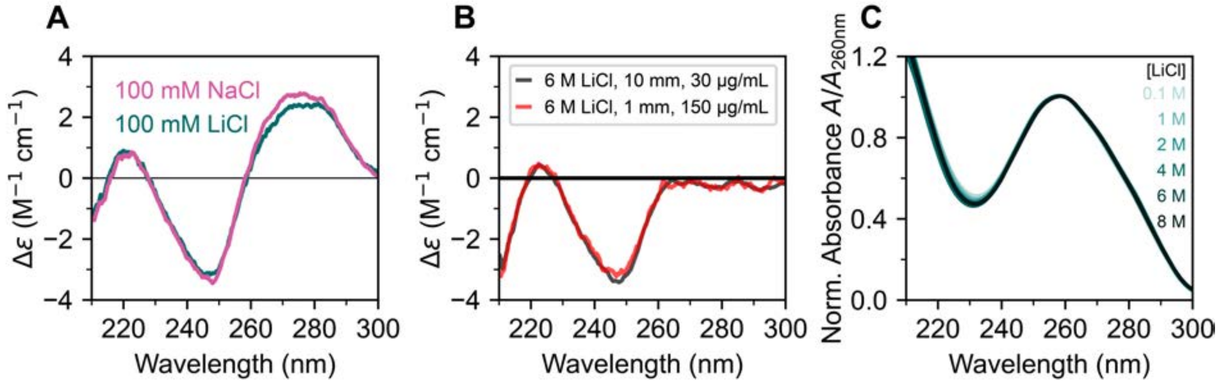


**Supplementary Figure S11. Circular Dichroism of DNA in varying salt concentrations.** A) CD spectra of λ-DNA in 100 mM NaCl and 100 mM LiCl. Slight ion specificity is observed as the magnitude of the band at 280 nm appears larger for NaCl than for LiCl. B) CD spectra of λ-DNA in 6 M LiCl solution for DNA concentrations of 30 µg/mL and 150 µg/mL (measured using 10 mm or 1 mm pathlength cuvettes, respectively, to achieve similar levels of signal-to-noise and to keep the absorption in a reasonable range). The resulting spectra are essentially within error, suggesting that intermolecular interactions do not significantly influence the CD spectra under our conditions. C) Normalized absorbance of λ-DNA for LiCl concentrations of 0.1-8 M. The spectra are essentially superimposable, indicating again the absence of formation of large DNA aggregates.

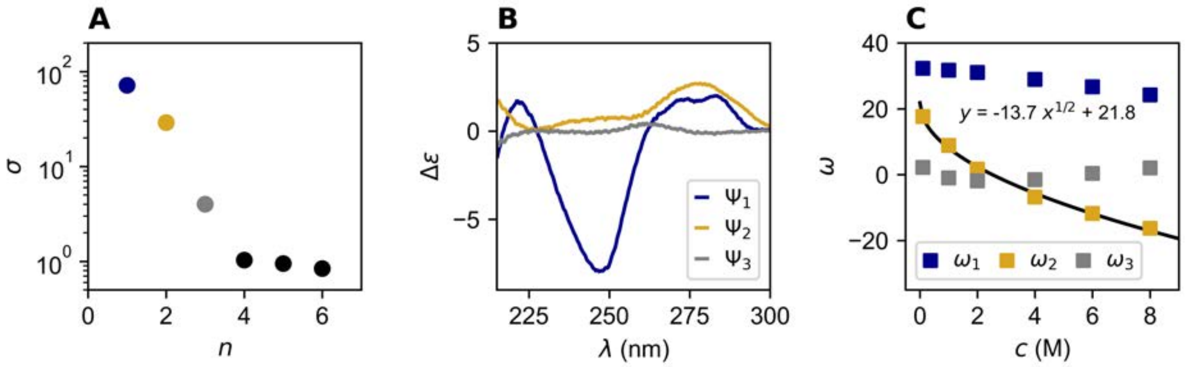


**Supplementary Figure S12. Singular value decomposition of CD spectra**. We applied analysis by singular value decomposition (SVD) to the CD spectra of λ-DNA (Figure 2) in varying concentrations of LiCl. SVD was implemented by representing the $i$-th observed CD spectrum $\{\overrightarrow{\Phi_i} | 1 \leq i \leq n\}$ of the DNA in salt condition $i$ as a vertical vector of dimension m measured for the wavelengths channels $\{\lambda_j | 1 \leq j \leq m\}$. The matrix M comprises all spectral vectors and is a m×n rectangular matrix where m > n.

The SVD transformation (13,14) decomposes the matrix of the CD spectra M as follows:

$$\mathrm{M} = \mathrm{UD}V^T = \mathrm{U}(\,\sigma_1\,\overrightarrow{v_1} \quad \sigma_2\,\overrightarrow{v_2}\,\ldots\,\sigma_r\,\overrightarrow{v_r}\,) = \mathrm{U}(\,\overrightarrow{\omega_1} \quad \overrightarrow{\omega_2}\,\ldots\,\overrightarrow{\omega_r}\,).$$

The matrix U consists of the basis function vectors Ψ as columns. The basis functions are not CD spectra themselves, but the spectra can be represented in the basis of the (orthogonal) $\Psi_i$. D is a diagonal n×n matrix that has the singular values $\sigma_i$ on the diagonal. The product $\mathrm{D}V^T = \vec{\omega}_i$ gives the (LiCl-dependent) weights of the basis functions $\Psi_i$ when constructing the scattering profiles from the basis of the $\Psi_i$.

A) Singular values $\sigma$ obtained from the singular value decomposition of the six experimentally determined CD spectra as a function of LiCl concentration. Singular values $\sigma$ indicate the relative importance of the basis functions for representing the data. While the first two components clearly have large singular values, the last three have small singular values and likely correspond to noise. The third singular value is somewhat intermediate. B) Basis functions Ψ for the three components with the highest $\sigma$ values. The first two components clearly contain signal while the third component is somewhat marginal. C) Coefficients corresponding to the basis functions $\omega$. The solid line is a fit of a square root dependance to the $\omega_2$ values.

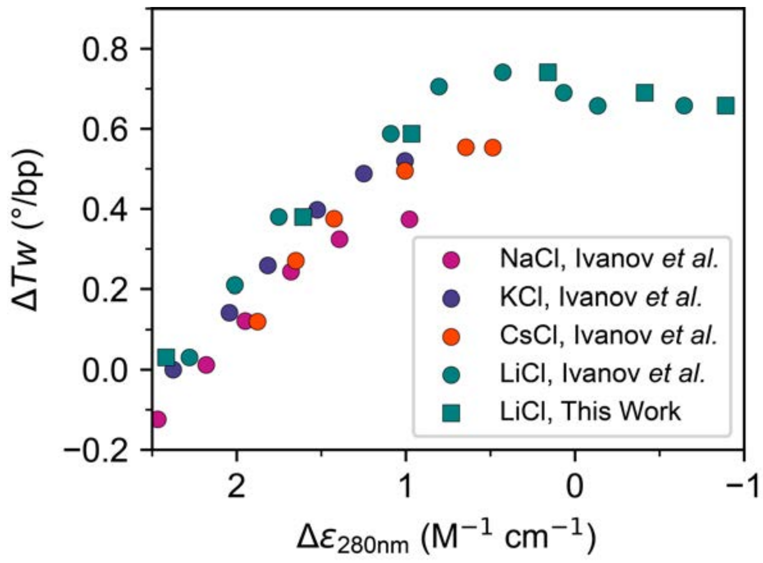


**Supplementary Figure S13. Changes in DNA twist vs. CD values at 280 nm for different monovalent salts.** The values for twist relative to the 100 mM KCl reference conditions were taken from the magnetic tweezers data obtained in this work (Table 1 and Figure 1G). Data for $\Delta\varepsilon_{280nm}$ were taken either from this work (green squares) or from the Ref. (15) (circles).

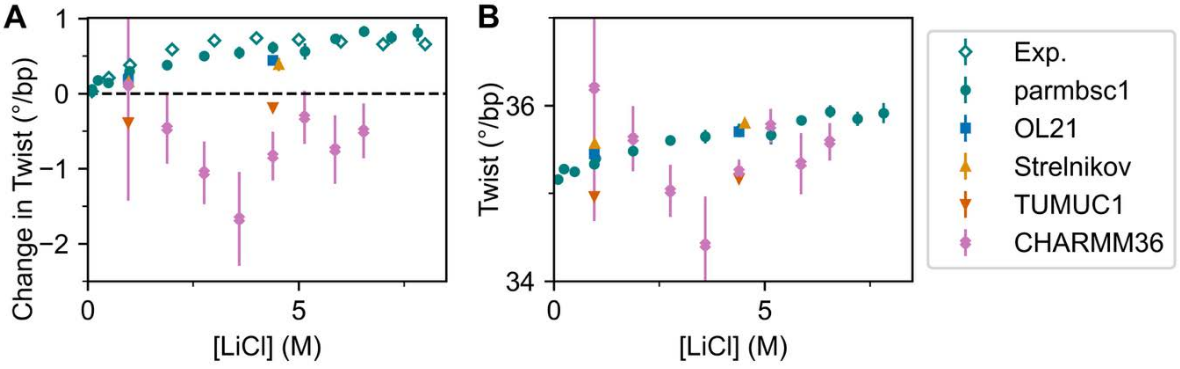


**Supplementary Figure S14. DNA twist from MD simulations for different force fields as a function of LiCl concentration.** A) Change in twist as a function of LiCl concentration relative to 0.10 M KCl from experiments and MD simulations with different force fields. A subset of the data are shown in Figure 3. The negative changes in twist from the simulations with TUMUC1 are caused by a too strong affinity of $Li^{+}$ towards the oxygen or nitrogen atoms of the nucleobases (**Supplementary Figures S15A and S18B**). The CHARMM36 simulations show very large scatter, due to the unzipping of the DNA ends (**Supplementary Figure S15B**). B) Absolute values of the DNA twist as a function of LiCl concentration. Data correspond to the mean ± SEM from six independent simulations.

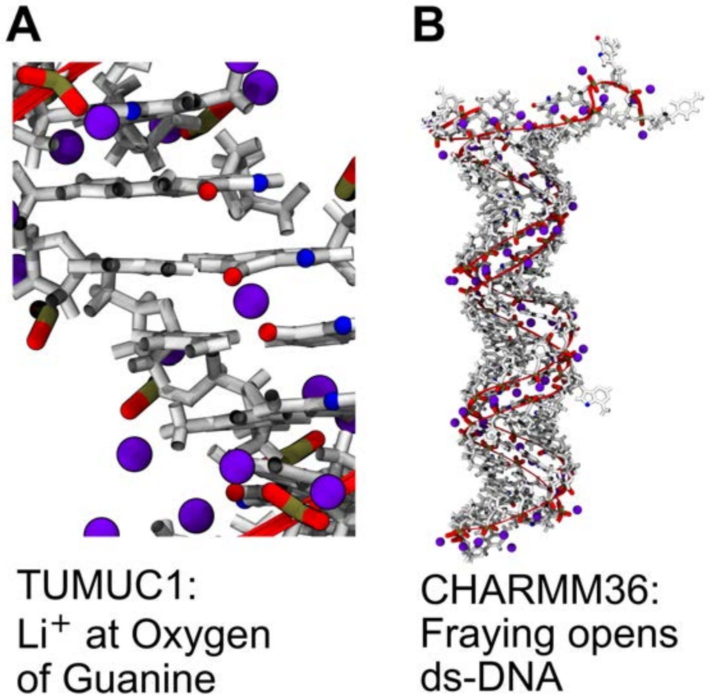


**Supplementary Figure S15. Simulation snapshots illustrating typical structures from simulations with the TUMUC1 and CHARMM36 force fields.** A) In case of TUMUC1, more $Li^+$ (purple) are adsorbed than with parmbsc1 (**Supplementary Figure S18)**. The strong adsorption involves the oxygen atoms of guanine, where the ions bridge stacked nucleobases. B) During the simulations with CHARMM36, the DNA ends unzip, resulting in unreliable values for the helical parameters. This opening is shown at the upper part of the depicted DNA structure, where one strand of the DNA backbone (red) is not paired with the other one.

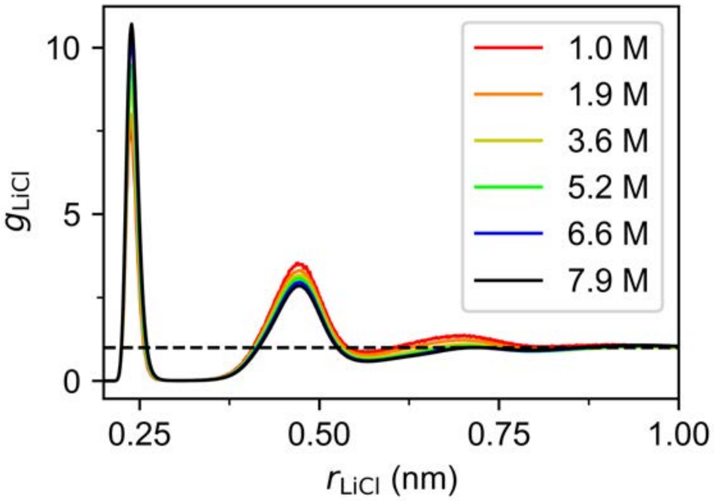


**Supplementary Figure S16. Radial distribution functions $g_{LiCl}$ along the distance $r_{LiCl}$ between $Li^+$ and $Cl^-$ for different concentrations of LiCl in a water box without DNA.** The black dashed line indicates unity, which is asymptotically reached at large distances. Simulations used the ion parameters from Mamatkulov-Schwierz in combination with the TIP3P water model. Note that the concentrations in the legend are reported in molarity and correspond to molalities of 1.0, 2.0, 4.0, 6.0, 8.0 and 10.0 m, respectively.

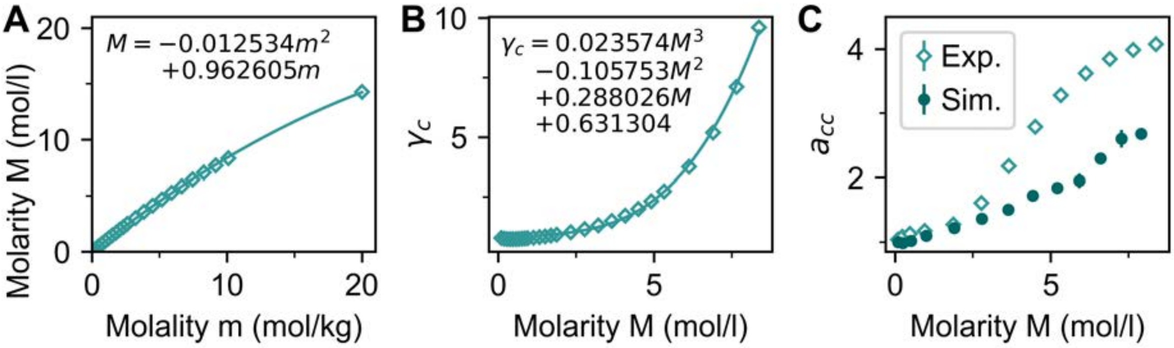


**Supplementary Figure S17. Activity coefficients and activity derivatives as function of LiCl concentration in water.** (A) Molarity plotted against molality for LiCl in water. Data are taken from Ref. (8). The line is a parabolic fit to the data. The equation for the fit is shown as an inset and used for the conversion from molality to molarity. (B) Activity coefficient $\gamma_c$ plotted against concentration (molarity). Points are a selection of values reported in (7). The solid line is a cubic fit to the data with coefficients shown in the inset. (C) Activity derivative $a_{cc}$ plotted against concentration. Shown are the results from experiments using numerical differentiation of the cubic fit to the data in panel B (open diamonds) and from MD simulations employing Kirkwood-Buff theory (filled circles). For the simulations, data points correspond to the mean ± SEM from three independent simulation runs.

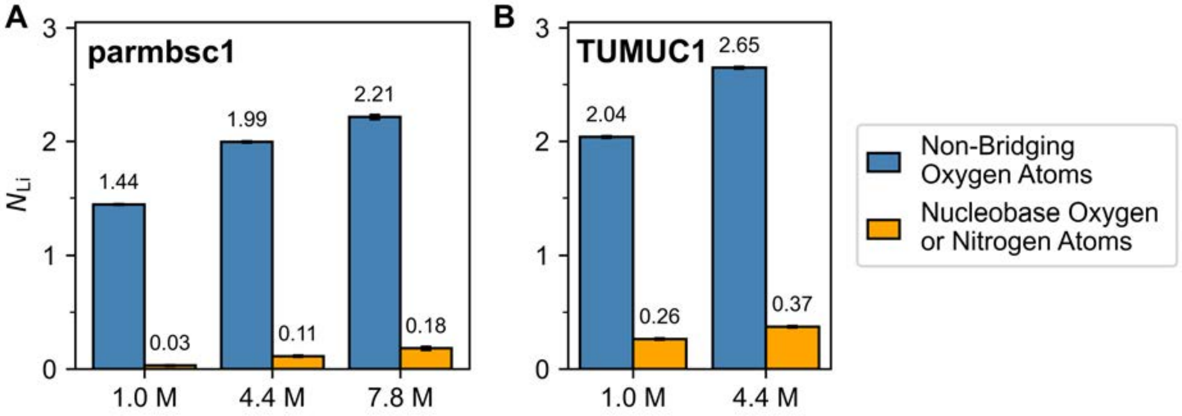


**Supplementary Figure S18. Ion adsorption with increasing LiCl concentration for two DNA force fields.** Average numbers of $Li^+$ ions per phosphate group $N_{\mathrm{Li}}$ that are within a distance of 0.3 nm to any non-bridging oxygen atom of the phosphate groups or nucleobase oxygen or nitrogen atoms from simulations (A) with the parmbsc1 force field or (B) with the TUMUC1 force field. Error bars show the standard error from six independent simulations. Note that the concentrations used as categories are reported in molarity and correspond to molalities of 1, 5, and 10 m, respectively.
